# Reovirus RNA recombination is sequence directed and generates internally deleted defective genome segments during passage

**DOI:** 10.1101/2020.10.19.346031

**Authors:** Sydni Caet Smith, Jennifer Gribble, Julia R. Diller, Michelle A. Wiebe, Timothy W. Thoner, Mark R. Denison, Kristen M. Ogden

## Abstract

For viruses with segmented genomes, genetic diversity is generated by genetic drift, reassortment, and recombination. Recombination produces RNA populations distinct from full-length gene segments and can influence viral population dynamics, persistence, and host immune responses. Viruses in the *Reoviridae* family, including rotavirus and mammalian orthoreovirus (reovirus), have been reported to package segments containing rearrangements or internal deletions. Rotaviruses with RNA segments containing rearrangements have been isolated from immunocompromised and immunocompetent children and *in vitro* following serial passage at high multiplicity. Reoviruses that package small, defective RNA segments have established chronic infections in cells and in mice. However, the mechanism and extent of *Reoviridae* RNA recombination are undefined. Towards filling this gap in knowledge, we determined the titers and RNA segment profiles for reovirus and rotavirus following serial passage in cultured cells. The viruses exhibited occasional titer reductions characteristic of interference. Reovirus strains frequently accumulated segments that retained 5′ and 3′ terminal sequences and featured large internal deletions, while similar segments were rarely detected in rotavirus populations. Using next-generation RNA-sequencing to analyze RNA molecules packaged in purified reovirus particles, we identified distinct recombination sites within individual viral gene segments. Recombination junction sites were frequently associated with short regions of identical sequence. Taken together, these findings suggest that reovirus accumulates defective gene segments featuring internal deletions during passage and undergoes sequence-directed recombination at distinct sites.

**IMPORTANCE:** Viruses in the *Reoviridae* family include important pathogens of humans and other animals and have segmented RNA genomes. Recombination in RNA virus populations can facilitate novel host exploration and increased disease severity. The extent, patterns, and mechanisms of *Reoviridae* recombination and the functions and effects of recombined RNA products are poorly understood. Here, we provide evidence that mammalian orthoreovirus regularly synthesizes RNA recombination products that retain terminal sequences but contain internal deletions, while rotavirus rarely synthesizes such products. Recombination occurs more frequently at specific sites in the mammalian orthoreovirus genome, and short regions of identical sequence are often detected at junction sites. These findings suggest that mammalian orthoreovirus recombination events are directed in part by RNA sequences. An improved understanding of recombined viral RNA synthesis may enhance our capacity to engineer improved vaccines and virotherapies in the future.

## INTRODUCTION

Genetic drift and reassortment are typically considered the primary mechanisms by which segmented RNA viruses acquire genetic diversity. However, recombination also occurs regularly during the replication of these viruses and yields non-canonical RNA molecules that differ from full-length genome segments (1, 2). These non-canonical RNAs may be packaged and can influence viral population dynamics, persistence, and host immune responses. A subset of non-canonical RNAs resulting from viral recombination events are known as defective viral genomes (DVGs), because they are unable to replicate in the absence of functional trans-complementation by the full-length parental RNA. The most frequently reported type of RNA DVG arises from recombination events resulting in large deletions that may remove much of the coding region of a viral RNA while retaining elements required for polymerase binding, replication, and packaging (2–4). Functions of DVGs are poorly defined for RNA viruses, although some configurations have demonstrated roles in viral replication interference, innate immune antagonism, and virus evolution (2). For segmented *Reoviridae* viruses, the need to successfully package a multi-partite genome likely imposes additional restrictions on the variety of non-canonical RNAs that are tolerated. Overall for the *Reoviridae,* there have been limited studies of recombination events and the recombined non-canonical RNAs they generate.

For the *Reoviridae* family of segmented, double-stranded (ds) RNA viruses, RNA segment termini are predicted to direct packaging and other viral replication processes. Rotavirus is the leading cause of diarrheal mortality among unvaccinated children under five years of age (5). Mammalian orthoreovirus (reovirus) has been linked to loss of oral tolerance associated with celiac disease and is in advanced clinical trials as an oncolytic (6). *Reoviridae* particles are non-enveloped, multi-layered, and encapsidate nine to twelve dsRNA genome segments (7). Following entry into target cells and outer capsid removal, subvirion particles function as nanoscale factories that transcribe capped, positive-sense viral RNA (+RNA) species that are ~0.7 to 4 kb in length (7–11). The ten reovirus segments are classified as large (L), medium (M), or small (S), while the eleven rotavirus segments are numbered. Most segments contain a single open reading frame (ORF) flanked by 5′ and 3′ untranslated regions (UTRs) (12, 13). Extensive base-pairing between 5′ and 3′ terminal regions, interrupted by secondary structures, has been predicted for many segments (14–17). *Reoviridae* +RNA sequences required for packaging, assortment, transcription, or translation encompass the UTR and extend into the ORF (17–21). Experimental and computational evidence suggests inter-segment complementarity drives assortment of *Reoviridae* +RNAs, with shorter gene segments nucleating +RNA complexes that are packaged into assembling viral capsids (22–26). Since *Reoviridae* viruses can package non-canonical segments, at least some level of divergence from the consensus sequence is tolerated (10, 25, 27–34). However, the extent of recombination and degree to which non-canonical segments are packaged are unknown.

For the *Reoviridae*, non-canonical gene segments have been reported. Rotaviruses with RNA segments containing rearrangements have been isolated from immunocompromised and immunocompetent children and *in vitro* following serial passage at high multiplicity (31, 33, 35). Most reported rearrangements involve partial head-to-tail duplications inserted after the functional ORF (35). In some cases, rotavirus rearrangements occur at preferred sites, with direct repeat sequences or RNA secondary structures proposed as recombination hot spots (28, 30, 33). Rearranged segments have also been detected in orbiviruses (29, 32). Finally, reoviruses establishing chronic infections *in vitro* and *in vivo* may package small, defective RNA segments (21, 27, 34, 36, 37). Together, these observations suggest that recombination is not a rare occurrence during *Reoviridae* replication. However, previous work often has not approached studies of *Reoviridae* recombination and non-canonical segments in a systematic manner. The mechanism of *Reoviridae* recombination, the frequency with which recombined RNAs are synthesized or packaged, and the functions of recombined RNAs remain poorly understood.

In the current study, we sought to elucidate the type and frequency of non-canonical RNA synthesis by *Reoviridae* viruses. We determined the titers and RNA segment profiles of reovirus (rsT1L and rsT3D^I^) and rotavirus (rsSA11) laboratory strains that had been rescued by reverse genetics then serially passaged ten times, each in triplicate lineages, in cultured cells. Viruses exhibited occasional reductions in titer that are characteristic of interference by defective-interfering viral genes. The two reoviruses accumulated non-canonical RNAs that retain 5′ and 3′ termini and feature one or more large internal deletions, while the rotavirus rarely accumulated such DVGs. Analyses of next-generation RNA-sequencing data sets from purified rsT1L reovirus RNA revealed many junctions, with hot spots for recombination in specific viral gene segments. Further, an overlap of 3-9 bp of identical sequence was favored at the recombination junction sites, suggesting that DVG formation is primarily determined by sequence complementarity. These findings suggest that reovirus frequently synthesizes and packages DVGs that contain internal deletions and undergoes recombination at distinct sites across the genome that encode key sequence features. This work provides rationale for future studies that will reveal detailed mechanisms of DVG synthesis and DVG effects on viral population dynamics and host responses.

## RESULTS

### Virus titer patterns differ among serially passaged *Reoviridae* viruses

To compare virus population replication patterns over multiple infections between reovirus and rotavirus in cultured cells, we utilized a serial passage approach with two strains of reovirus, T1L and T3D^I^, and one strain of rotavirus, SA11. T1L and T3D represent distinct human reovirus serotypes (12). T3D induces necrosis and substantially more apoptosis than T1L in murine L929 fibroblasts (L cells) (38, 39). Recombinant strain (rs) T3D^I^ is identical to T3D, except for a T249I mutation in attachment protein σ1 that renders it resistant to proteolysis (19). SA11 is a simian rotavirus strain that induces necrosis in African green monkey kidney epithelial (MA104) cells (40). For each passage series, the viruses were recovered using plasmid-based reverse genetics, and working virus stocks were made after one additional round of amplification in cultured cells (Fig. 1A). rsT1L, rsT3D^I^, and rsSA11 were serially passaged ten times, each in three lineages, and infectious virus titers were determined. We found that while the three lineages of each virus strain exhibited relatively similar virus titer patterns, rsT1L, rsT3D^I^, and rsSA11 exhibited notable differences in virus titer pattern across the passage series (Fig. 1B-D). Virus titers for rsT1L reovirus climbed steadily from ~ 3 × 10^5^ PFU/ml at passage 1 (P1) to ~ 10^12^ PFU/ml at P5, remained high through P7, decreased by ~10-fold to 1000-fold at P8, then rebounded through P10 (Fig. 1B). For serially passaged rsT3D^I^ reovirus, virus titers climbed from ~ 3 × 10^5^ PFU/ml at P1 to ~ 6 × 10^7^ PFU/ml at P5, then fluctuated between ~ 3 × 10^7^ and 4 × 10^8^ PFU/ml through P10, with some variability observed among lineages (Fig. 1C). By P1, rsSA11 rotavirus had already reached titers of ~5 × 10^7^ PFU/ml (Fig. 1D). Titers then decreased by ~5-fold to 25-fold at P3 and P9, with a rebound to peak titers approaching 10^8^ PFU/ml during intervening passages. Thus, titer patterns in serially passaged rsSA11 rotavirus lineages resembled those of rsT3D^I^ reovirus in that peak titers were lower and stayed within a narrower range, but they resembled those of rsT1L reovirus in that distinct titer dips and rebounds were observed concurrently for all three lineages (Fig. 1B-D).

**Figure 1.**
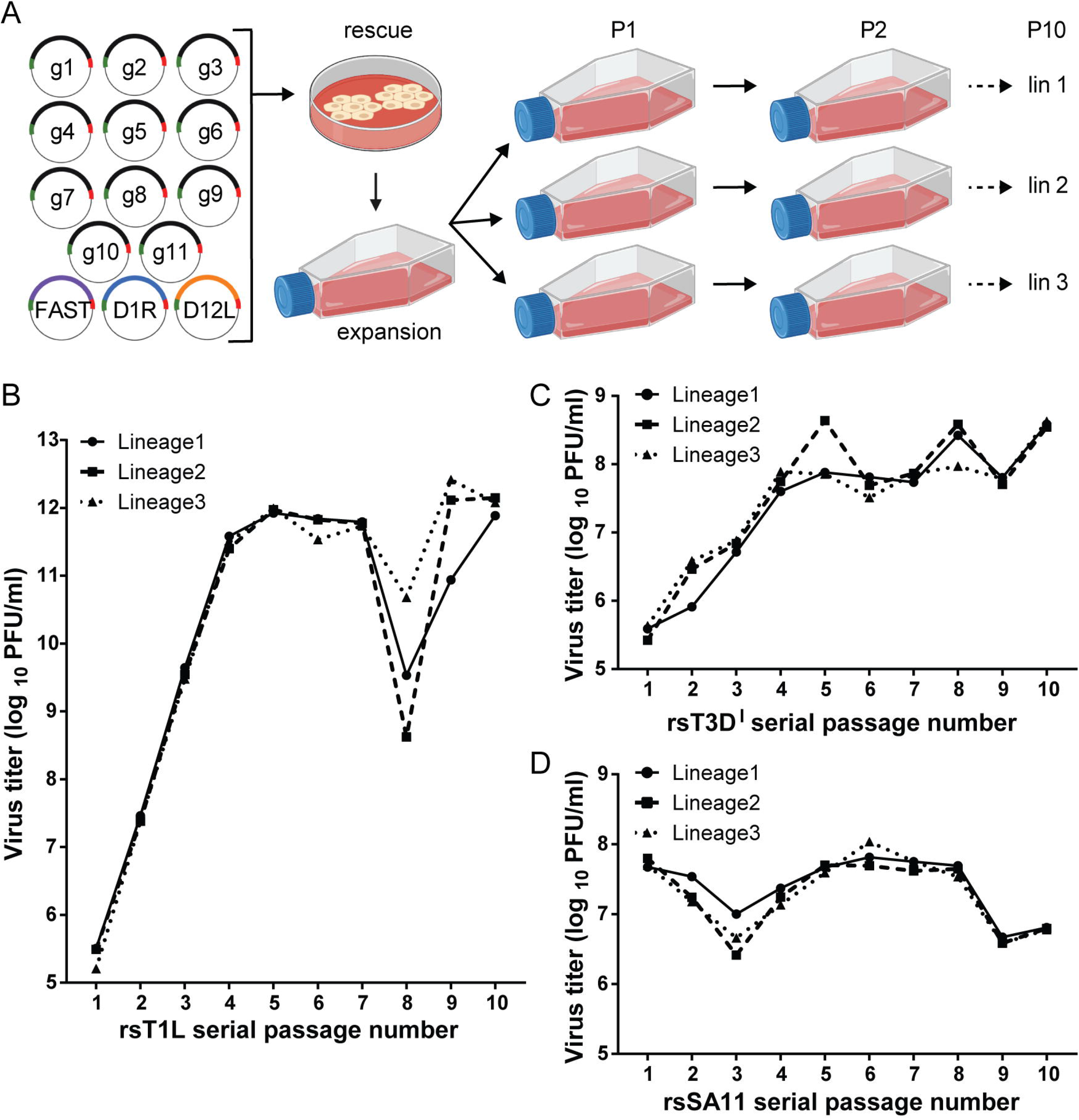
Serial passage workflow and virus titers. (A) Workflow for rotavirus serial passaging. Baby hamster kidney cells expressing T7 RNA polymerase were transfected with plasmids encoding the eleven rotavirus +RNAs (g1-g11) and helper plasmids encoding capping enzymes (D1R and D12L) and a cell-cell fusion protein (FAST) then co-cultured with MA104 cells to promote recombinant virus rescue. rsSA11 was amplified by a single passage in MA104 cells, and virus stock titer was determined. To generate P1 stocks, MA104 monolayers in three flasks were adsorbed at a MOI of 0.25 PFU/cell, washed, and incubated with fresh medium for 48 h prior to lysis by multiple rounds of freezing and thawing. Subsequent passages (P2-P10) were generated by adsorption of MA104 monolayers with three milliliters of cleared lysate from the previous passage, with three lineages each passaged in an independent series. A similar workflow was used for rescue and passaging of reovirus, employing standard reverse genetics approaches (19, 56) and with passages conducted in suspension rather than monolayer culture. (B-C) Graphs showing titers for three lineages of rsT1L (B) or rsT3D^I^ (C) reoviruses across ten serial passages, quantified by plaque assay. (D) Graphs showing titers for three lineages of rsSA11 rotavirus across all ten serial passages, quantified by fluorescent focus assay.

### Reovirus and rotavirus produce small non-canonical segments during serial passage

Alterations to virus titer patterns during serial passage can be due to the presence of DVGs, which may interfere with viral replication (4, 41). To detect the presence of non-canonical segments in lysates from reovirus and rotavirus passage series, we used reverse transcription polymerase chain reaction (RT-PCR). RNA molecules containing both the 5′ and 3′ termini of a specific viral gene segment were amplified and visualized (Table 1). Using this approach, we consistently detected full-length viral gene segments ≤ 3.3 kb in length, which included M and S reovirus segments and all rotavirus segments (g1-g11) (Figs. 2–3). Longer full-length segments, such as reovirus L segments, were amplified infrequently, due to limitations of the enzymes used for amplification. Thus, our assay conditions permitted detection of small RNA molecules that contained native viral 5′ and 3′ segment termini. For rsT1L reovirus lineage 1 (lin1), RT-PCR products smaller than the full-length segment were amplified from multiple passages for each segment, except S2 (Fig. 2A). Non-canonical segments derived from segments L1, M1, and S1 were detected beginning at about P2 and continuously for the remaining passages. For segments L2, L3, M2, and M3, non-canonical segments were detected only transiently. For segments M1, S1, and S4, an RNA product slightly longer than the full-length genome segment was detected. There is no non-canonical segment whose presence or absence clearly correlates with the changes in titer shown in Fig. 1B. However, a non-canonical L3 segment was detected from P5-P7, preceding the sharp decrease in titer at P8, and a faint, transient, non-canonical M1 segment was detected only in P8 and P9 (Figs. 1B and 2A).

**Table 1.**
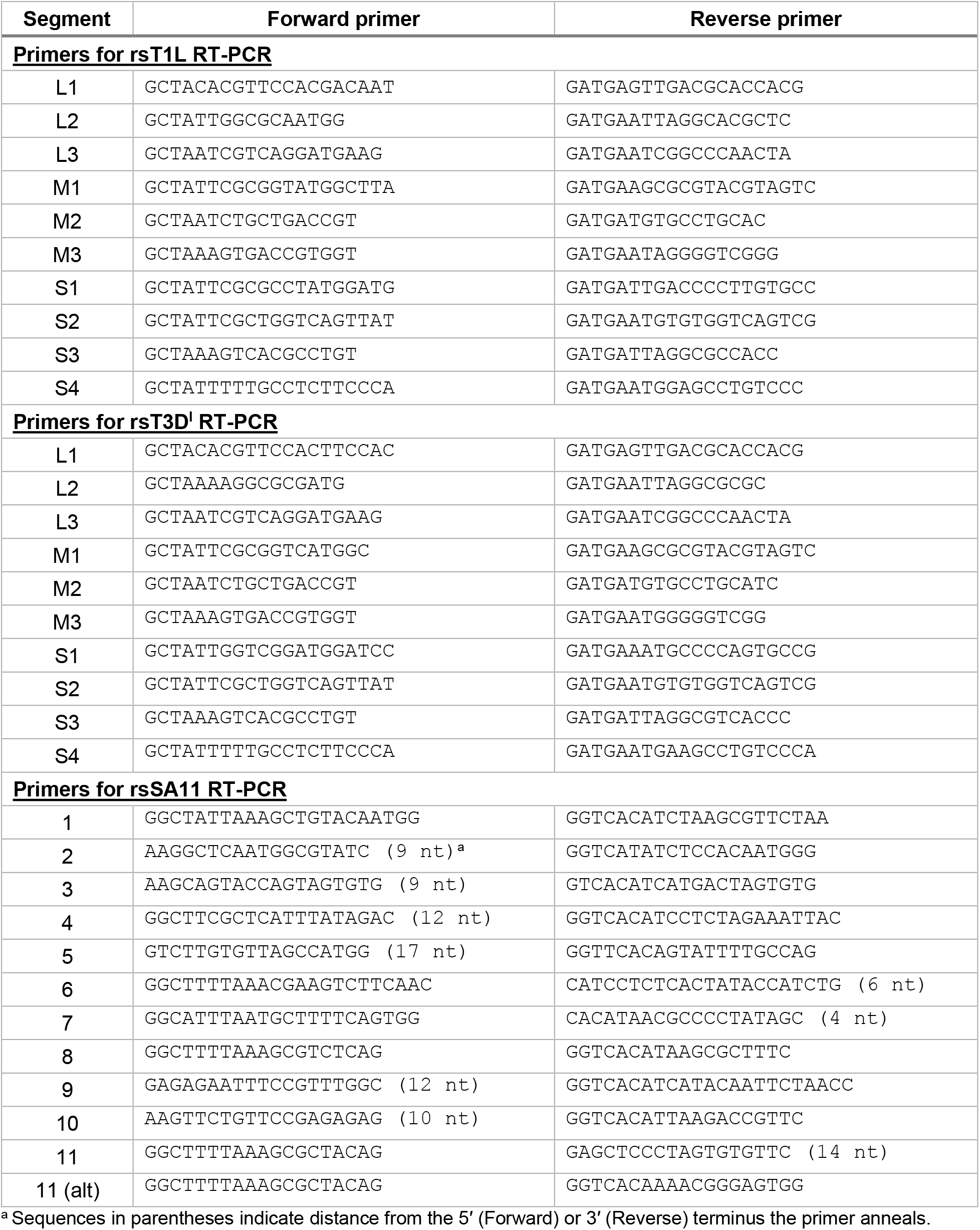
RT-PCR primers used in the current study.

**Figure 2.**
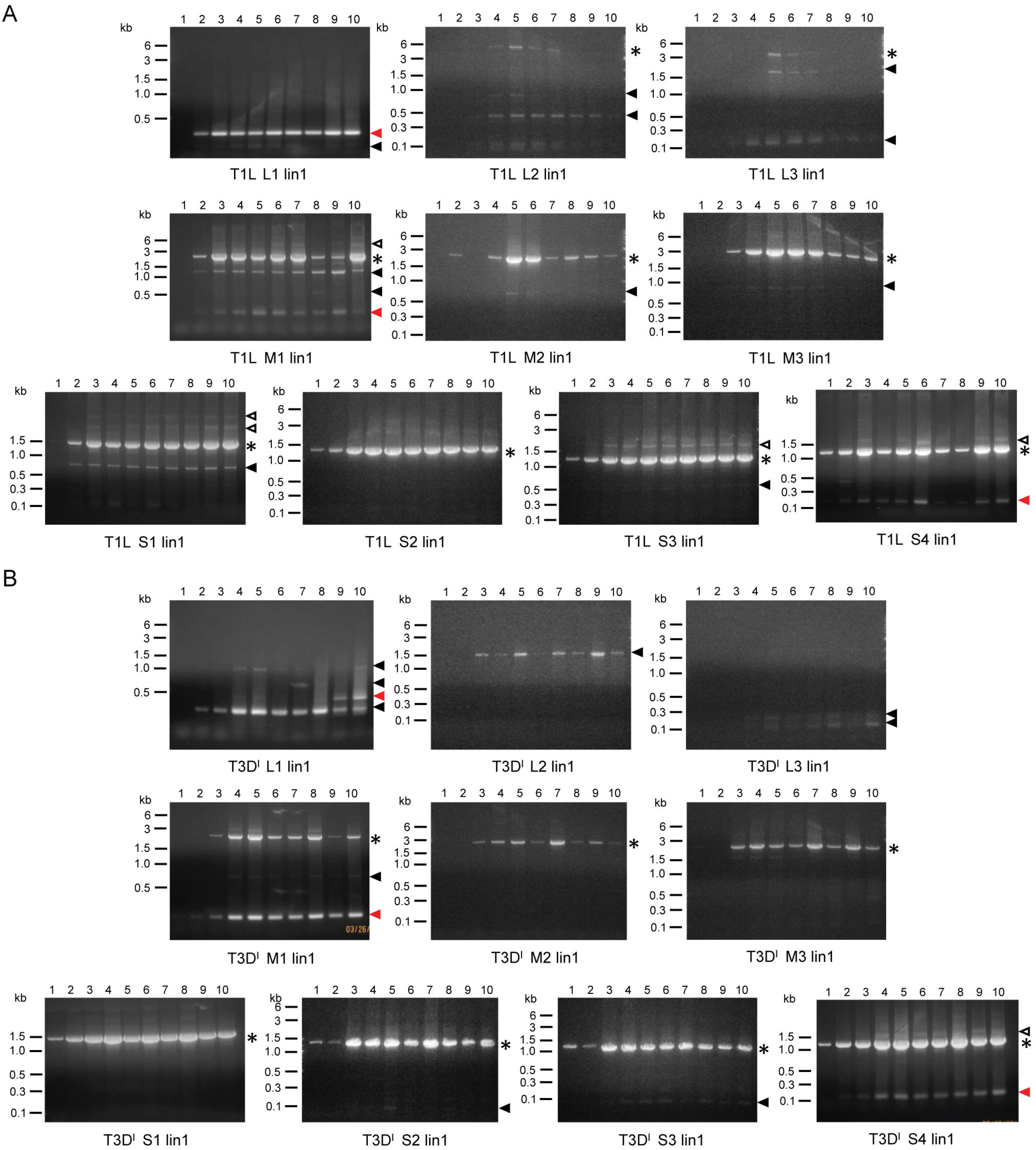
Serial passage lineage 1 reovirus segment profiles. RNA extracted from serial passage lysates was used as template in RT-PCR reactions, along with primers that bind to the 3’ and 5’ terminal sequences of each reovirus gene segment. Products from reactions in which template RNA was extracted from rsT1L lin1 P1-10 (A) or rsT3D^I^ lin1 P1-10 (B) were resolved on 1.2% ethidium-bromide stained agarose gels. The identity of the reovirus segment (L1-L3, M1-M3, S1-S4) recognized by the terminal primers is indicated. Black asterisks indicate the position of the full-length segment. Filled triangles indicate non-canonical RNA products smaller than the full-length segment. Red triangles indicate products that were excised and sequenced. Open triangles indicate products larger than the full-length segment.

**Figure 3.**
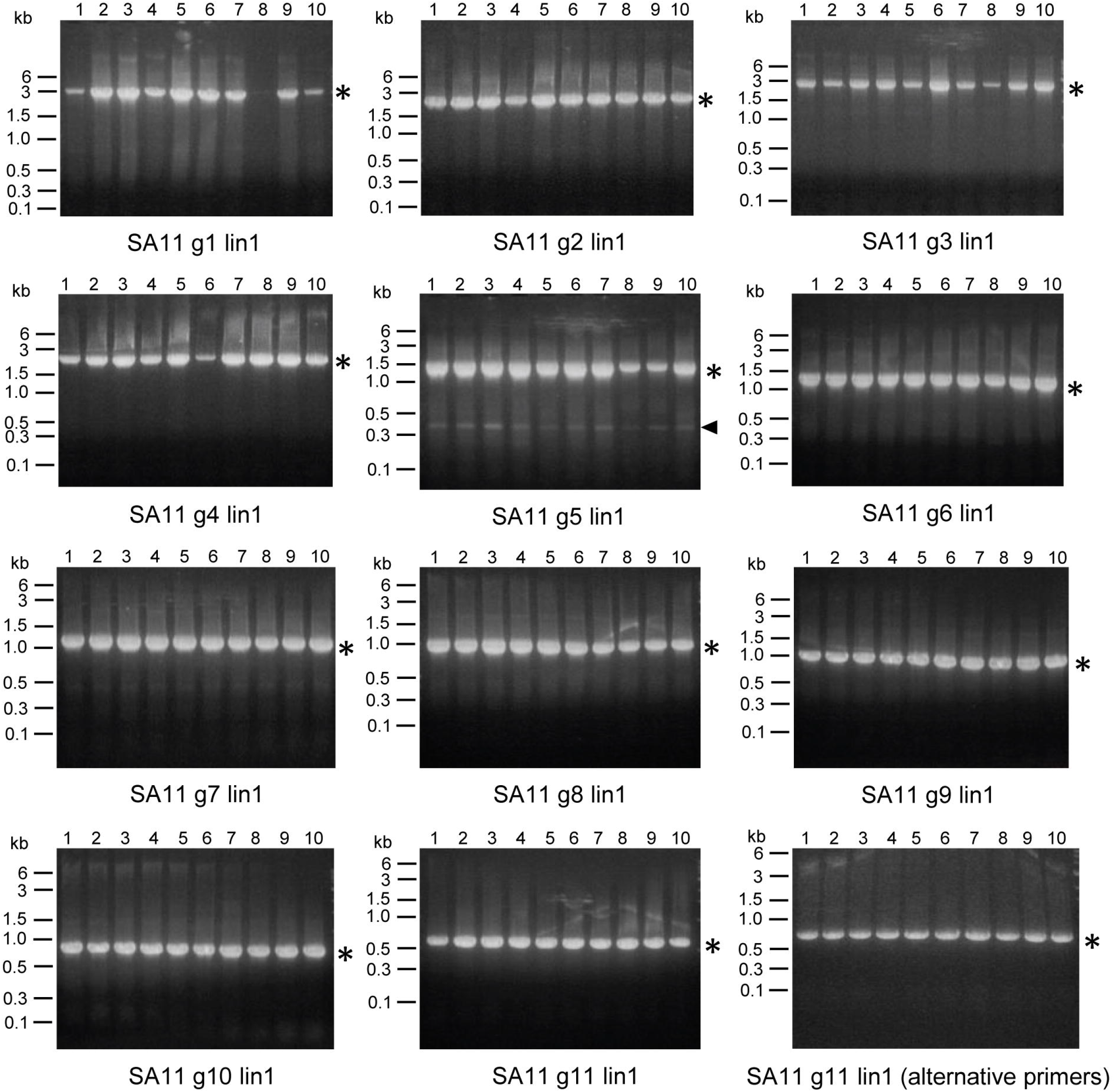
Serial passage lineage 1 rotavirus segment profiles. RNA extracted from serial passage lysates was used as template in RT-PCR reactions, along with primers that bind to the 3’ and 5’ terminal sequences of each rotavirus gene segment. Products from reactions in which template RNA was extracted from rsSA11 lin1 P1-10 were resolved on 1.2% ethidium-bromide stained agarose gels. The identity of the rotavirus segment (g1-g11) recognized by the terminal primers is indicated. Black asterisks indicate the position of the full-length segment. Filled triangles indicate products smaller than the full-length segment.

For rsT3D^I^ reovirus lin1, RT-PCR products smaller than the full-length segment were amplified from multiple passages for seven of ten segments (Fig. 2B). Like the patterns observed for rsT1L, these products often were detected by P3 and then continuously across the majority of rsT3D^I^ passages. However, some products were detected transiently. Products derived from the rsT3D^I^ L1 segment provide the most striking example of this property. The smallest non-canonical L1 segment was detected in P2-P10. A non-canonical L1 segment of ~ 1kb was detected strongly in P4-P5 and P10 but weakly in some other passages, and two intermediately sized non-canonical L1 segments were detected only in P7 or only in P9-P10. For segment S4, a non-canonical segment was detected that is slightly longer than the full-length genome segment.

In contrast to the frequent detection of non-canonical segments smaller than the full-length segment for serially passaged reoviruses, such a product was detected for only one of eleven rotavirus segments, g5, using primers that bind near the termini (Fig. 3). This non-canonical segment was present across all serial passages. For several segments, indistinct bands high on the gels suggested the presence of RT-PCR products that were longer than the full-length products. To determine whether the failure to detect small non-canonical segments for rsSA11 rotavirus was due to primer bias, we repeated the RT-PCR for g11, for which the initial primer bound 14 nucleotides internal to the 3′ end, using an extreme terminal primer (Table 1, alt primer). Using the new g11 primer pair, we still failed to detect any RT-PCR products smaller than the full-length segment (Fig. 3). The segment for which we had detected a small non-canonical segment was g5, and the 5′ primer bound a position beginning 17 nucleotides internal to the terminus. Taken together, lin1 RNA profiles suggest non-canonical RNA species that differ in length but have identical termini to the parental segments accumulate variably during reovirus and rotavirus serial passage in cultured cells.

After examining RNA profiles for all ten or eleven segments of lin1, we compared RNA profiles among all three lineages for each virus passage series following amplification by RT-PCR and resolution in 1.2% agarose, as in Figs. 2–3. For rsT1L and rsT3D^I^ reoviruses, we used primers that anneal to the termini of L1, M1, and S4, and for rsSA11 rotavirus, we used primers that anneal to g2, g5, and g11 (Table 1). These segments were chosen to compare RNA profiles between representative small, medium, and large viral gene segments. For rsT1L reovirus, we found that profiles of putative DVGs were mostly conserved across the three independently passaged lineages (Fig. 4A). For rsT1L L1, in all three lineages, a single, bright band of ~0.3 kb was detected beginning at P2 or P3 and continuing through P10, and a smaller, faint band was detected in some central passages. For rsT1L M1, non-canonical RNA segments were detected at ~1.5 kb and ~ 0.2 kb in several passages. For rsT1L S4, non-canonical RNA segments just slightly larger than the full-length segment and at ~0.15 kb were detected in several passages.

**Figure 4.**
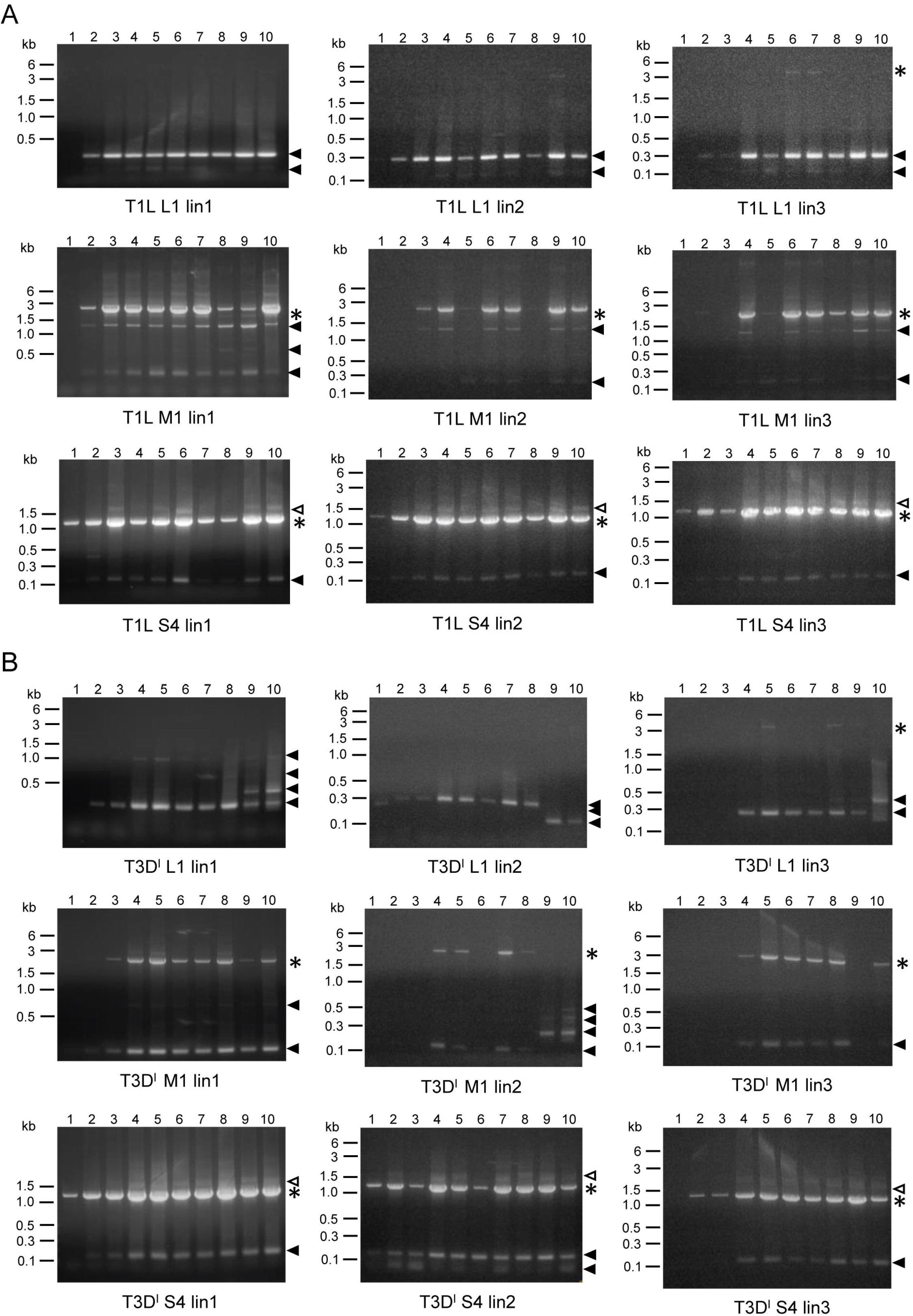
Serial passage lineage 1-3 reovirus segment profiles. RNA extracted from serial passage lysates was used as template in RT-PCR reactions, along with primers that bind to the 3’ and 5’ terminal sequences of reovirus gene segments L1, M1, or S4. Products from reactions in which template RNA was extracted from rsT1L reovirus lin2-3 P1-10 (A) or rsT3D^I^ reovirus lin2-3 P1-10 (B) were resolved on 1.2% ethidium-bromide stained agarose gels. Gels from lin1 are identical to those shown in Figure 2 but are included for comparison. Black asterisks indicate the position of the full-length segment. Filled triangles indicate products smaller than the full-length segment. Open triangles indicate products larger than the full-length segment.

In contrast to rsT1L reovirus RNA profiles, those of rsT3D^I^ reovirus were less well conserved across the three lineages (Fig. 4B). For rsT3D^I^ L1, all three lineages contained a small non-canonical RNA segment, ~0.3 kb in size, present in many passages. However, lin1 contained transient, larger non-canonical RNA segments, as described above. Lin2 contained smaller, transient rsT3D^I^ L1 non-canonical RNA segments instead of the ~0.3 kb segment in P1, P9, and P10. Lin3 contained a larger non-canonical rsT3D^I^ L1 segment in later passages. For rsT3D^I^ M1, many passages of all three lineages contained a small putative non-canonical RNA segments of ~0.1 kb, but this segment appeared absent from the last two passages of lin2 and lin3, and additional larger non-canonical M1 segments were detected in P9-P10 for lin2. The rsT3D^I^ S4 RNA profiles were more consistent among lineages than those of the M1 and L1 segments and were similar to those of rsT1L S4, with non-canonical RNA segments just slightly larger than the full-length segment and at ~0.15 kb detected in several passages (Fig. 4). For rsT3D^I^ lin2, an additional non-canonical S4 segment of < 0.1 kb also was detected in several passages.

RNA profiles for rsSA11 rotavirus were very similar across the three lineages, featuring a single non-canonical RNA segment of ~0.4 kb only for g5 (Fig. 5). As noted above for lin1, for some segments in lin2 and lin3, indistinct bands high on the gels suggested the presence of RT-PCR products that were longer than the full-length products. The observation that similarly sized products that differ from full-length segments were amplified by primers specific to the 5′ and 3′ segment termini arose in multiple independent passages of rsT1L reovirus and rsSA11 rotavirus suggests that these viruses preferentially accumulate certain non-canonical RNAs.

**Figure 5.**
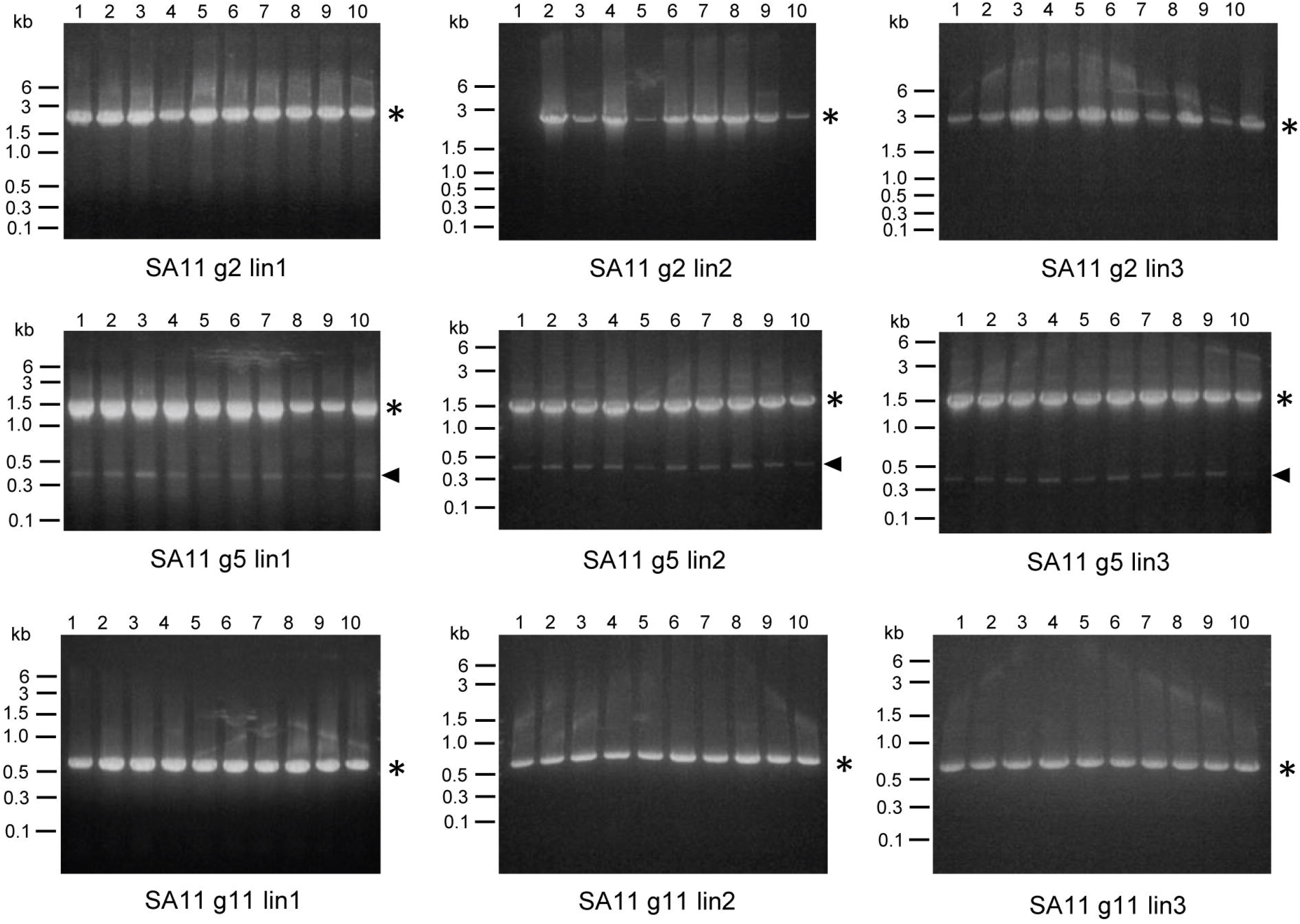
Serial passage lineage 1-3 rotavirus segment profiles. RNA extracted from serial passage lysates was used as template in RT-PCR reactions, along with primers that bind to the 3’ and 5’ terminal sequences of rotavirus gene segments g2, g5, or g11. Products from reactions in which template RNA was extracted from rsSA11 rotavirus lin2-3 P1-10 were resolved on 1.2% ethidium-bromide stained agarose gels. Gels from lin1 are identical to those shown in Figure 3 but are included for comparison. Black asterisks indicate the position of the full-length segment. Filled triangles indicate products smaller than the full-length segment.

### Non-canonical reovirus segments feature large deletions

To gain insight into the identities of non-canonical RNA products detected in lysates of serially passaged reovirus, we excised bands from agarose gels on which RT-PCR products had been resolved (Fig. 2) and used the Sanger method to determine their sequences. We selected bright, low-molecular weight bands amplified using primers specific for the termini of rsT1L and rsT3D reovirus L1, M1, and S4 (Table. 1). The sequenced segments ranged from ~ 7% to 33% of the length of the parental segments. Comparison with reference sequences revealed that each product was a DVG that contained relatively short intact termini and one or two large internal deletions (Fig. 6). The largest single deletion, in the rsT1L L1 DVG, was of 3,581 nucleotides and removed the majority of the ORF and about half of the 3′ UTR. In every case, the 5′ UTR was left intact, and for five of the six sequenced DVGs, more than 50 nucleotides of the 5′ end of the ORF also was left intact (Fig. S1). The shortest terminal regions on either side of a novel DVG junction were only 15-20 nucleotides in length and were observed at the 5′ terminus of the rsT3D^I^ M1 DVG and the 3′ termini of the rsT1L L1 DVG and the rsT3D^I^ S4 DVG. For the rsT1L L1 DVG and the rsT3D^I^ S4 DVG, part of the 3′ UTR was deleted during its emergence. For the rsT1L M1 DVG, less than 50 nucleotides of the ORF preceding the 3′ UTR was retained, but at 80 nucleotides this UTR is relatively long. Together, these observations suggest that serially passaged reoviruses regularly undergo recombination events resulting in generation of DVGs that retain 5’ and 3’ termini and feature one or multiple large internal deletions.

**Figure 6.**
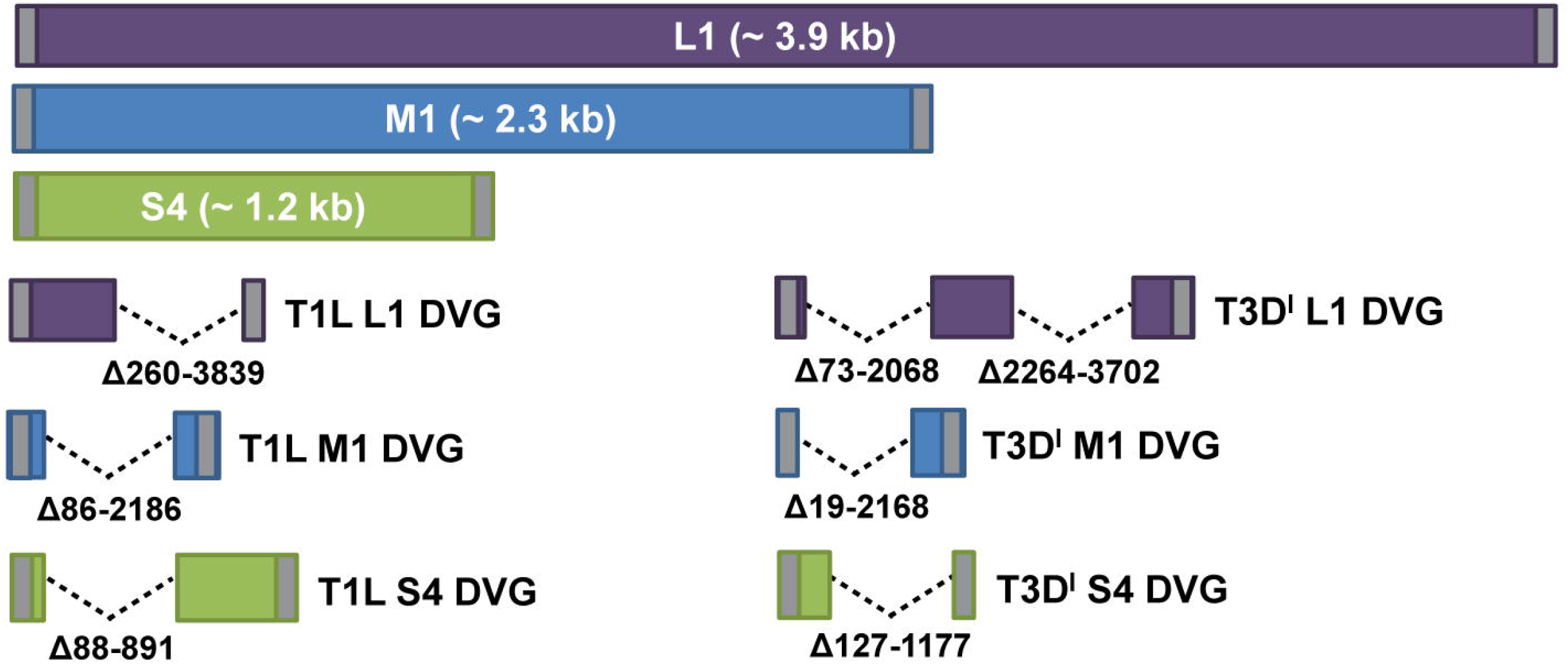
Schematics of reovirus DVGs. RT-PCR products amplified using primers that bind the 5′ and 3′ termini of the L1, M1, and S4 reovirus segments, smaller than the full-length segments, and indicated in Figure 2 were excised from agarose gels and sequenced. Schematics, drawn to scale, of reovirus L1, M1, and S4 segments and sequenced rsT1L and rsT3D^I^ reovirus DVGs are shown. The ORF is colored purple (L1), blue (M1), or green (S4), and the 5′ and 3′ UTRs are colored gray. Deletions are indicated by dotted lines, and deleted nucleotides are described.

To better understand reovirus recombination, we examined sequences immediately surrounding the new junctions created in DVGs (Table 2). In several cases, a pattern was elusive. However, for the rsT1L L1 DVG, the first rsT3D^I^ L1 DVG junction, and the rsT3D^I^ S4 DVG, short regions of sequence similarity (six to seven nucleotides) were detected downstream from the recombination site. For the rsT1L L1 DVG, the upstream four nucleotides also were identical. Thus, these junctions were identical before and after recombination, or, in the case of the rsT1L L1 DVG, contained a single inserted nucleotide.

**Table 2.**
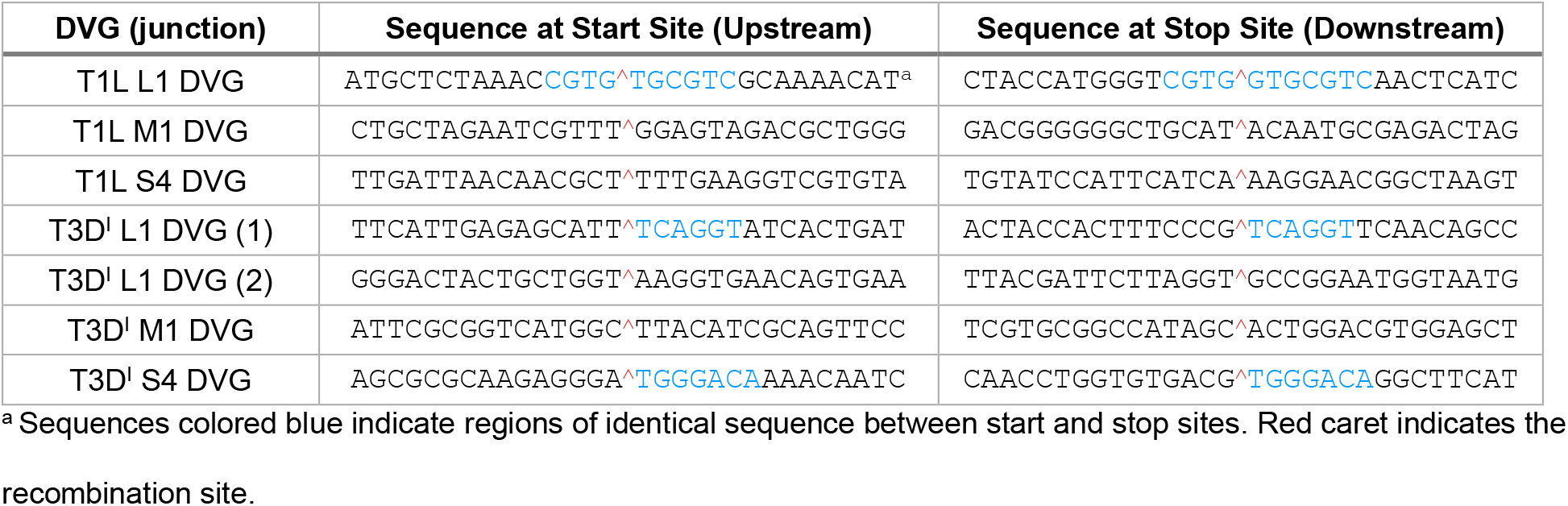
Nucleotides surrounding sequenced reovirus DVG recombination junctions.

### Reovirus recombination occurs at distinct sites

After noting the short regions of identical sequence immediately downstream or spanning the junction sites for some of the DVGs we excised and sequenced, we wondered whether analysis of larger numbers of junction sites would reveal reovirus recombination biases Therefore, we extracted RNA from benzonase-treated (to remove extra-virion nucleic acids) rsT1L particles, generated randomly primed, directional libraries, and sequenced them using Illumina technology. These viruses had been amplified only a few times following recovery using plasmid-based reverse genetics to make laboratory stocks and were not anticipated to have accumulated numerous DVGs. We aligned high-quality reads to reference rsT1L sequences for each individual gene segment using *ViReMa* (*Vi*rus *Re*combination *Ma*pper), a platform that detects recombination events in viral genomes from input next-generation sequencing data, and analyzed recombination events with a custom bioinformatic pipeline (42, 43). Our sequencing averaged ~10^4^ reads per site across most genome segments, with reduced coverage at the extreme 5′ and 3′ termini (Figs. 7A and S2). Approximately 99% of reads mapped to viral gene segments, demonstrating that the virion particles had been sufficiently purified (Table 3). We detected a genome-wide recombination frequency of approximately 0.01% and identified many non-canonical junctions within each viral gene segment (Figs. 7B and S3, Table 3). In accordance with previous reports, a recombination junction was defined as a deletion greater than 5 base-pairs flanked both upstream and downstream by a 25 base-pair high-quality alignment (42, 43). Forward, 5’ to 3’ junctions were filtered for each segment. Small, internal deletions of less than 150 bp with an average size of ~66 bp were especially prevalent in the L segments, as indicated by the concentration of points along the diagonal axes of junction plots in Figures 7B and S3, though similar deletions were detected in all segments. The S segments, particularly S2, more frequently exhibited fusion of one segment terminus to the other than did the larger segments, as indicated by the concentration of points in the lower right corners of junction plots in Figures 7B and S3. Several segments exhibited hot spots for recombination (Figs. 7C and S4). A hot spot was defined as a position or clustering of positions where the recombination frequency exceeded that of the overall genome. Further, positions of interest were identified if the recombination frequency was higher than that of the rest of the segment. For example, in segment M2, recombination frequencies at nucleotides 953 and 2049 were at least four times higher than those at most other positions across the segment. In segment S4, hot spots were detected around nucleotides 117-131, 404-452, 506-532, 921, and 999-1005. However, for a few segments (L2, S2, and S3), recombination frequency was low across the entire gene segment length (Fig. S4). Thus, the identification of positions in the reovirus gene segments where the recombination frequency was up to 100-fold higher than the global recombination frequency demonstrates that reovirus recombination may occur at key, distinct sites across gene segments.

**Table 3.**
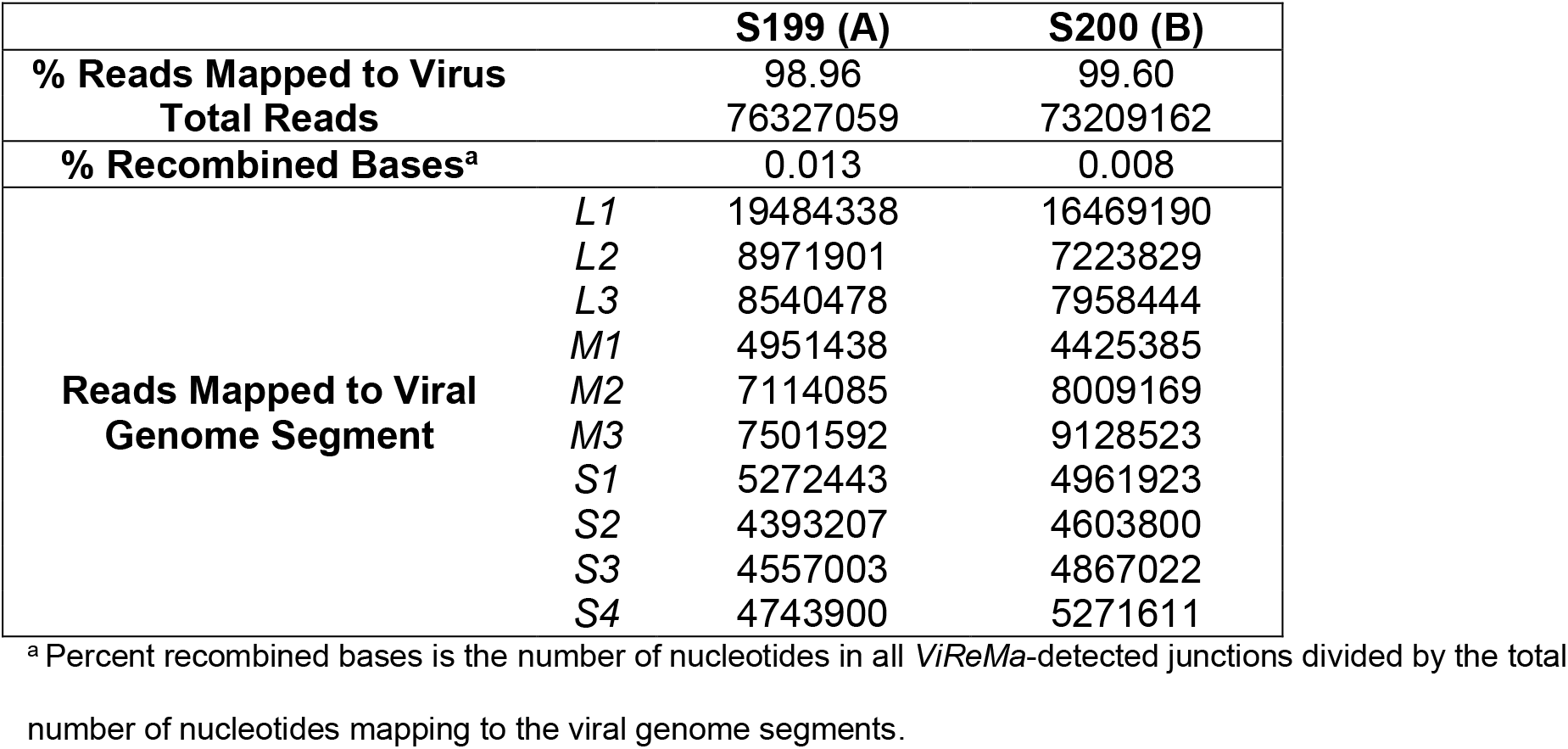
Alignment statistics.

|  |  | S199 (A) | S200 (B) |
| --- | --- | --- | --- |
| % Reads Mapped to Virus |  | 98.96 | 99.60 |
| Total Reads |  | 76327059 | 73209162 |
| % Recombined Bases<sup>a</sup> |  | 0.013 | 0.008 |
| Reads Mapped to Viral Genome Segment | L1 | 19484338 | 16469190 |
|  | L2 | 8971901 | 7223829 |
|  | L3 | 8540478 | 7958444 |
|  | M1 | 4951438 | 4425385 |
|  | M2 | 7114085 | 8009169 |
|  | M3 | 7501592 | 9128523 |
|  | S1 | 5272443 | 4961923 |
|  | S2 | 4393207 | 4603800 |
|  | S3 | 4557003 | 4867022 |
|  | S4 | 4743900 | 5271611 |

<sup>a</sup> Percent recombined bases is the number of nucleotides in all *ViReMa*-detected junctions divided by the total number of nucleotides mapping to the viral genome segments.

**Figure 7.**
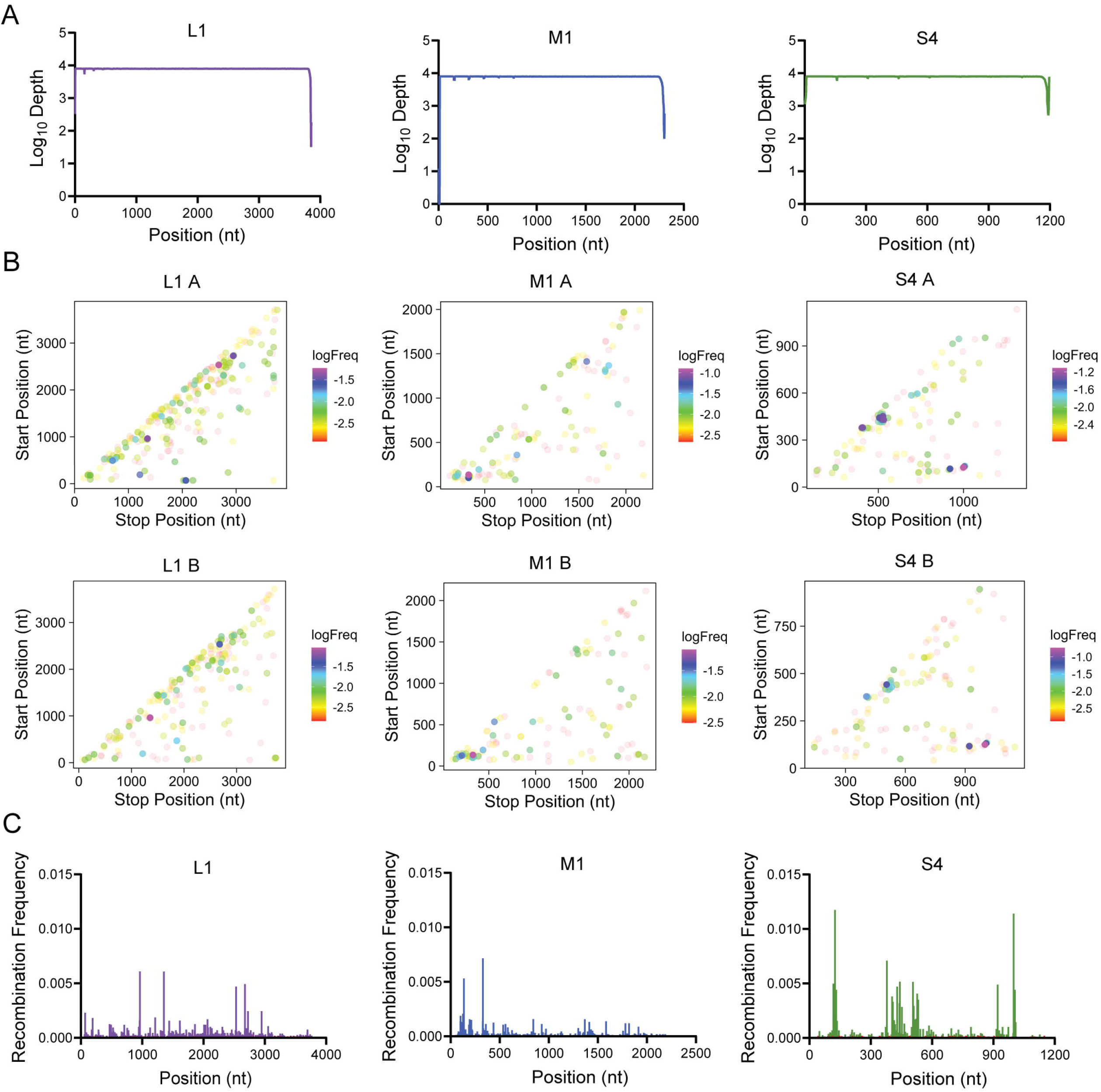
rsT1L reovirus junction frequency. (A) Graphs showing sequence depth at each nucleotide position across the indicated rsT1L genome segment. Mean coverage depth for two samples of sequenced virion RNA is shown. (B) Graphs showing recombination junction site location and frequency in sequenced rsT1L virion RNA for the indicated reovirus genome segment. Junction sites are indicated by dots whose position corresponds to upstream and downstream sequences that are merged to form a novel junction. Junction frequency is indicated by dot color, according to the legend to the right of each image. Junction maps for independently purified and sequenced virion RNA preparations are shown in separate images, labeled A or B. (C) Graphs showing recombination frequency at each nucleotide position across the indicated rsT1L genome segment. Mean positional recombination frequency for two samples of sequenced virion RNA is shown.

To test whether reovirus DVGs encode specific sequence preference, we extracted and quantified the upstream and downstream sequences flanking both the junction start and stop sites (Fig 8A). The percent adenosine (A), cytosine (C), guanosine (G), and uracil (U) was calculated and plotted for each position in a 10-nt window flanking the junction site represented by a red line and caret (^) (Figs 8B and S5). To exclude potential biases for overrepresented species, junctions were not weighted by depth. The global composition including all segments was compared to the L1, M1, and S4 gene segments individually (Fig. 8B). There was not a strong enrichment or depletion of any nucleotide across all segments. Similarly, the L1, M1, and S4 segments did not display strong nucleotide preferences with a few notable exceptions. The M1 junction start sites were enriched for cytosine. Both the start and stop sites of S4 were relatively enriched for adenosine immediately upstream of the junction sites. Finally, both the start and stop sites of L1 were relatively enriched for guanosine immediately upstream of the junction. Thus, while the reovirus junction sites did not demonstrate a strong overall sequence preference for a particular nucleotide or sequence motif, recombination at sites with specific nucleotide compositions may be vary in a segment-dependent manner.

**Figure 8.**
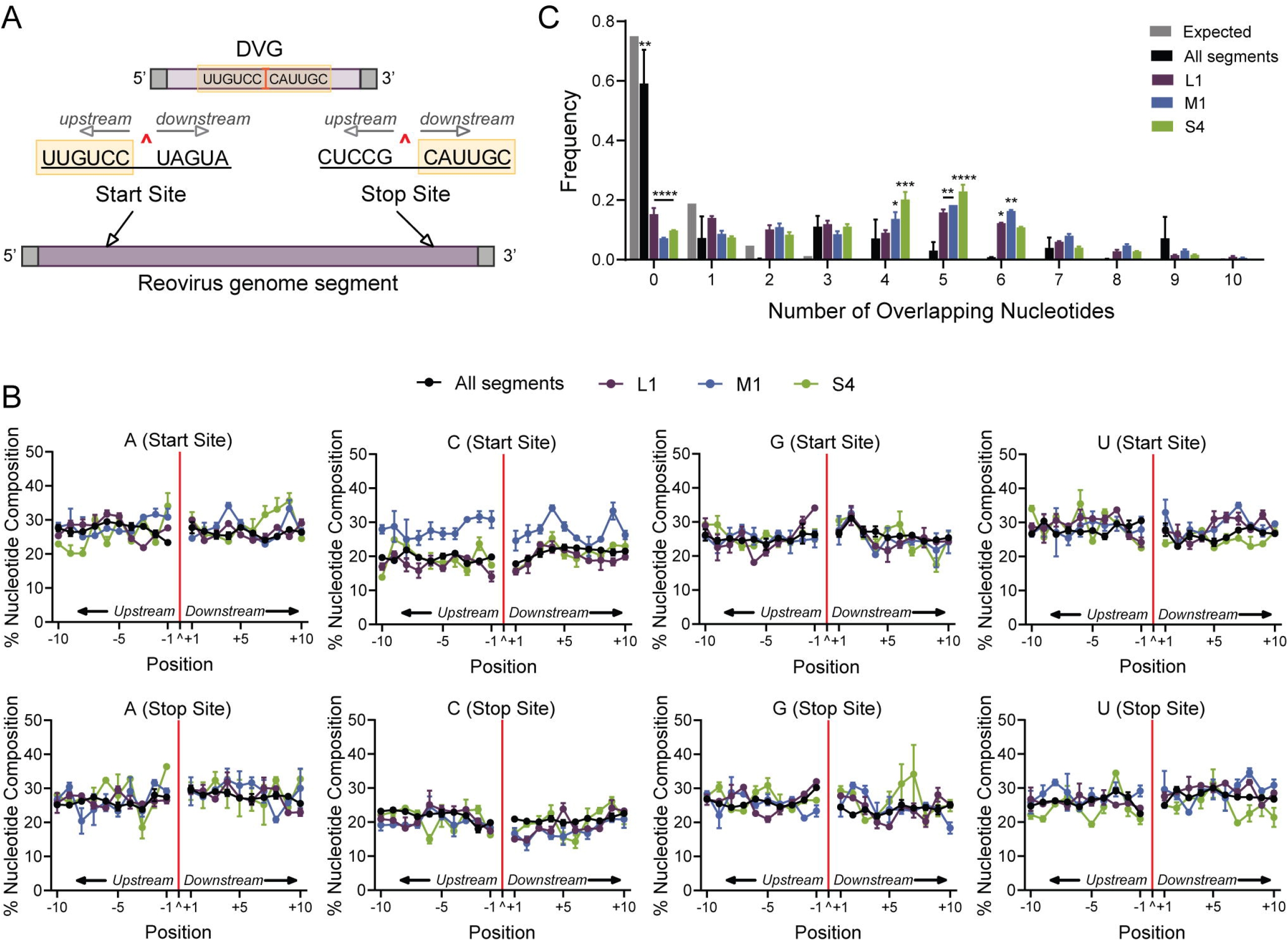
rsT1L reovirus junction site composition. Nucleotide composition and sequence homology were quantified from junction start and stop sites. (A) Schematic of DVG recombination junctions in the context of the parental gene segment. The junction is labeled with a red caret (^) and example sequences are shown. DVG junction sites are characterized by a 5′ start site and a 3′ stop site. Sequences upstream of the start site and downstream of the stop site (orange highlight) form the junction-spanning sequence of the DVG. (B) Nucleotide composition was calculated as the percent adenosine (A), cytosine (C), guanine (G), and uracil (U) at each position in a 10-base pair region surrounding the DVG start and stop sites. The junction is labeled as a caret (^) and denoted with a solid red line. Positions upstream (−10 to −1) and downstream (+1 to +10) of the junction position are indicated. Each point represents a mean (*n* = 2) and error bars represent standard error. (C) Sequence microhomology distributions of all rsT1L gene segments (black) and the L1 (purple), M1 (blue), and S4 (green) gene segments were compared to an expected probability distribution (gray). The frequency of each overlap is displayed as a mean (*n* = 2), and error bars represent standard error.

We further tested whether reovirus recombination occurs in regions of sequence microhomology, similar to other RNA viruses, such as dengue virus and flock house virus (FHV) (44). Microhomology was defined as a region of identical sequence between 2 and 20 base pairs (bp) in length. When microhomology at *ViReMa*-detected junctions was compared to an expected probability distribution, there was significant enrichment for 4-6 bp of overlap at junction sites for segments L1, M1, and S4 (Fig 8C). To ascertain any preference for sequence homology in the most frequently detected recombination junctions, we examined sequences upstream and downstream from the start and stop sites of the 25 most frequently identified S4 recombination junctions (Table 4). At least four nucleotides of identical sequence were detected for all but one of the top-ranking recombination junctions (Table 4). Typically, regions of sequence similarity spanned the recombination junction, such that sequences immediately surrounding the newly recombined junction site were identical to those at both the start and stop junction sites. Together, these observations suggest that RNA sequence homology plays a role in reovirus recombination site selection.

**Table 4.**
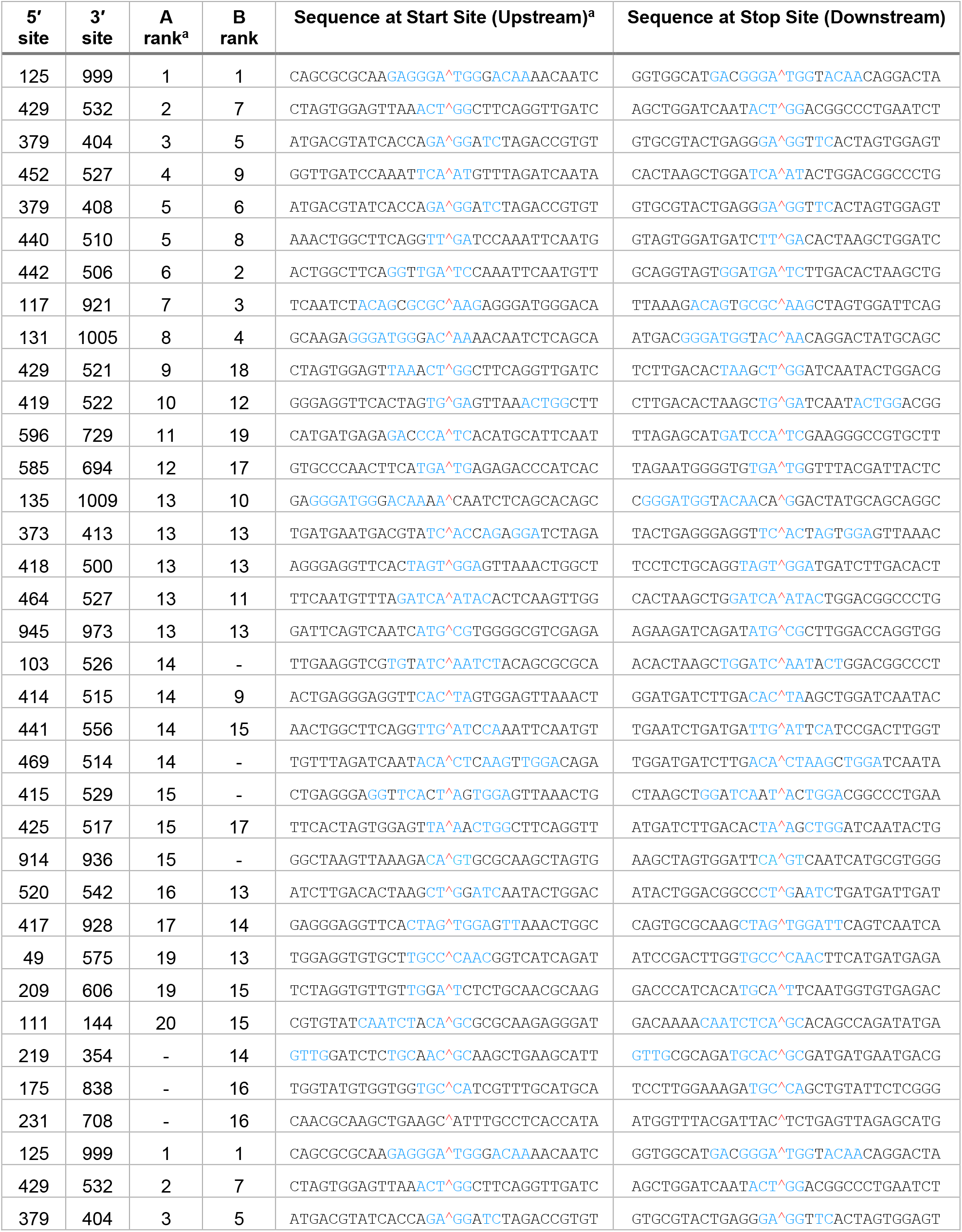
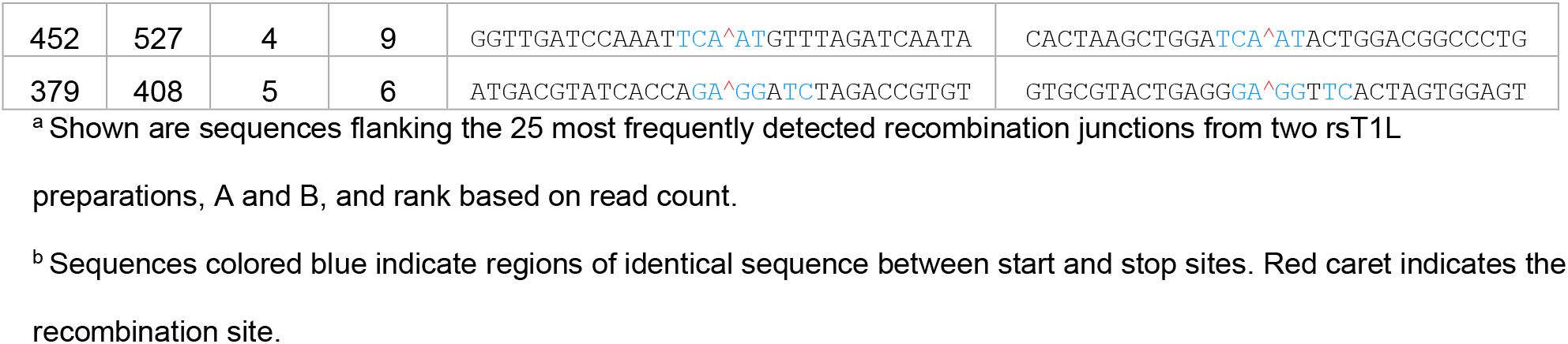
Nucleotides surrounding rsT1L S4 recombination junctions.

## DISCUSSION

While the synthesis and packaging of non-canonical segments, including DVGs, has been reported for multi-segmented, dsRNA viruses of the *Reoviridae* family (21, 28–37), the frequency and mechanism of recombination yielding these RNA products are unknown. Here, we serially passaged two strains of reovirus and one strain of rotavirus in cultured cells and compared the virus titers and RNA segment profiles. We demonstrated that small, non-canonical RNA products that retain the termini and feature large internal deletions are synthesized frequently by both strains of reovirus (Figs. 2, 4, and 6). Some small reovirus DVGs were maintained throughout the passage series, while other DVGs were transient (Figs. 2 and 4). RNA-seq analysis of RNA extracted from two independent preparations of purified, low-passage, benzonase-treated rsT1L reovirus revealed specific junction locations and recombination hot spots in individual gene segments (Figs. 7 and S3-S4). Sequences surrounding the recombination junction sites in packaged rsT1L RNA revealed significant nucleotide overlap and regions of sequence microhomology at junction sites for multiple gene segments (Fig. 8C and Table 4). Together, these findings suggest that reovirus frequently synthesizes and packages defective gene segments featuring internal deletions and that recombination can be directed by sequence and occurs at distinct sites.

Multiple factors may influence virus titer during serial passage. In the current study, virus titer patterns for serially passaged rsT1L and rsSA11 were similar in that distinct titer dips and rebounds were observed concurrently for all three lineages, but titer patterns for passaged rsT3D^I^ and rsSA11 lineages were similar in that peak titers were lower and stayed within a narrower range (Fig. 1B-D). T3D reovirus induces necrosis and substantially more apoptosis than T1L in L cells, and SA11 induces necrosis in MA104 cells (38–40). Once most cells in the culture have become infected, cell death may limit the peak titer that can be achieved for rsT3D^I^ and rsSA11, which may in part explain their narrower titer ranges. A repeating pattern of dips and rebounds in titer has been reported for serially and continuously cultured viruses and associated with generation of defective defective interfering particles (4, 27, 41, 45, 46). DVGs may interfere with viral replication through competition with full-length genomes for the viral polymerase and structural proteins (47). Small DVGs may be transcribed or replicated with higher efficiency than the full-length genome, permitting eventual monopolization of these resources. Particles encapsidating defective genomes can also potentily activate innate immune responses, which may interfere with viral replication (4). For all three lineages, rsT1L reovirus exhibited a noteworthy decrease in titer at P8, and rsSA11 rotavirus exhibited dips in titer at P3 and P9, each followed by a rebound (Fig. 1B and D). In contrast, rsT3D^I^ reovirus exhibited smaller fluctuations in titer that were not always parallel among lineages (Fig. 1C). The frequency of the titer dips and rebounds of rsT3D^I^ is more consistent with that previously reported for serially passaged bovine rotavirus than the frequency exhibited by the other viruses (45). Additional experiments are needed before any causal relationship between DVG presence or absence and virus titer changes can be established for the viruses in this study. However, some small DVGs were maintained throughout the passage series, suggesting they have no negative impact on the virus population, while other DVGs were transient (Figs.2–5). Viruses with the most consistent titer patterns among lineages, rsT1L and rsSA11, also had the most consistent RT-PCR profiles, while the virus with the most variability in titer among lineages, rsT3D^I^, also exhibited the most diverse RNA segment profiles, suggesting a potential link between these phenotypes (Figs. 1B-D and 4–5). We anticipate that the true diversity and abundance of DVGs present in reovirus and rotavirus populations is much greater than revealed by RT-PCR and electrophoretic analysis, which likely was limited to the most abundant populations of small DVGs with termini that match those of a specific gene segment.

Differences in DVG profiles among *Reoviridae* genera and reovirus strains detected in this study suggest underlying differences in recombination properties. It is unclear why serially passaged rsSA11 rotavirus lysates contained so few DVGs with matched segment termini and internal deletions, while such DVGs were detected for most rT1L and rsT3D^I^ reovirus segments in most passages (Figs. 2–5). For both reovirus and rotavirus, we detected putative non-canonical segments of greater length than the parental segment. In the literature, head-to-tail duplications (concatemers) resulting in longer segments have been most often reported for rotavirus, whereas internal deletions have been reported for reoviruses (21, 35, 36). It is possible that *Reoviridae* virion structure influences non-canonical RNA length. Reoviruses have turrets at the icosahedral vertices, through which newly synthesized viral +RNAs exit the particle, whereas rotaviruses are non-turreted *Reoviridae* family members. Like rotavirus, non-turreted orbiviruses package concatemers that are significantly longer than full-length segments (29). While mechanisms underlying differences in non-canonical viral segment synthesis remain unknown, infrequent detection of small rotavirus DVGs and frequent detection of small reovirus DVGs in this study (Figs. 2–5) are consistent with published findings.

It is unclear why rsT1L reovirus exhibited similar DVG profiles across lineages, while rsT3D^I^ exhibited much greater variability (Fig. 4). Consistency in RT-PCR profiles among rsT1L lineages and the finding that junction sites in packaged rsT1L RNA from two different low-passage virus preparations were extremely well conserved support a model in which rsT1L recombination occurs at distinct sites and yields specific DVGs (Figs. 4, 7B, and S3). Alternatively, rsT1L replication may tolerate DVG diversity poorly and limit the range of detected variants through purifying selection. The variability in RNA segment profiles among rsT3D^I^ lineages indicates that the products are not random artifacts of detection and suggests reduced recombination specificity or increased recombination frequency when compared to rsT1L (Fig. 4). For influenza virus, a segmented negative-strand RNA virus, the abundance of DVGs can be significantly affected by specific amino acid substitutions in the viral RNA-dependent RNA polymerase (RdRp), suggesting an important role for this enzyme in DVG synthesis (48, 49). For reovirus, the viral RdRp also likely plays a prominent role in recombination potential, with other viral and host factors potentially influencing recombination properties. For example, viral RdRp co-factor μ2 has been shown to differentially influence T1L and T3D transcriptional activity and replication efficiency in specific cell types (50, 51). Thus, identifying key determinants of recombination and non-canonical segment synthesis will be important areas of future study.

In comparison to other recent reports on RNA virus recombination, reovirus demonstrated ~37-fold lower recombination frequency than coronavirus and ~20-fold lower recombination frequency than FHV (43, 52). While microhomology can occur as an artifact of Illumina RNA-seq library preparation, artifactual recombination typically joins fragments in opposite orientations, involves short deletions, and lacks site specificity across the genome, whereas biological recombination by the virus often will join fragments in the same orientation and may occur at hot spots in the genome (42, 44, 52). Potential artifacts were removed by filtering for forward recombination junctions and removing short deletions from datasets presented here. Thus, despite the low reovirus recombination frequency detected in this study, the bioinformatic quality controls, orientation of joined fragments, and presence hot spots at specific sites in the genome support the biological relevance for recombination detected by RNA-seq (Figs. 7 and S3-S4). Detection and sequencing of DVGs containing large internal deletions following reovirus passage in cultured cells further supports the validity of these findings (Figs. 2, 4, and 6). Recombination frequency for FHV increases during passage and also may do so for passaged reoviruses (52).

The molecular mechanism of reovirus recombination is unknown. For influenza virus, DVGs containing internal deletions are common, and the 5′ and 3′ sequences of a DVG are always derived from the same segment and polarity (53). While our methodology cannot exclude the presence of other types of DVGs, our deep-sequencing data and previously reported findings suggest reovirus synthesizes DVGs that feature internal deletions and 5′ and 3′ termini from a single segment of the same polarity (Fig. 6) (21). Influenza DVGs containing internal deletions are proposed to arise from pausing of the RdRp during nascent strand synthesis while continuing to process along the template molecule and then resuming synthesis at a downstream point on the template (1). Downstream RdRp reinitiation may be guided by complementarity between the nascent strand and the site of reinitiation on the template (54, 55). Given the similarity in DVG composition and conservation of sequences at junction sites detected for multiple segments (Figs. 6 and 8C and Table 4), reovirus may synthesize DVGs using a mechanism similar to that proposed for influenza virus. RT-PCR products differing in size from full-length rsT1L S2, which exhibited low recombination frequency across the segment, were not detected (Figs. 2A and S4). However, such RT-PCR products were detected for L2 and S3, which also exhibited low overall recombination frequencies. Aside from the presence of small stretches of homologous sequence, it is unclear why recombination occurs more frequently at some sites than others throughout the rsT1L reovirus genome (Figs. 7C and S4). It is possible that, in addition to sequence, +RNA secondary structure also contributes to recombination site selection.

Effects of non-canonical RNA species, including DVGs, during *Reoviridae* infection are poorly understood. In addition to their effects on viral populations and the host, studies of non-canonical segments will reveal principles governing assortment and packaging, which may enable more sophisticated engineering of *Reoviridae*-based vaccines and therapeutics using reverse genetics platforms. While passaging at relatively high MOI and detection by RT-PCR suggests the existence of small DVGs in reovirus populations, these approaches are relatively insensitive (Figs. 1–6). The frequent detection of novel junctions resulting from recombination events via next-generation sequencing in relatively low-passage reovirus stocks suggests these events are common even in the absence of high-passage conditions (Figs. 7–8 and S2-S5).

Continued application of sensitive approaches will enable a deeper understanding of recombination frequency, and the identities of non-canonical RNA species, mechanisms by which they are generated, and their impacts on virus populations and the host.

## MATERIALS AND METHODS

### Cells

Spinner-adapted L929 cells were grown in suspension culture in Joklik’s minimum essential medium (JMEM; US Biological) supplemented to contain 5% fetal bovine serum (FBS) (Gibco), and 2 mM L-glutamine (Corning). All media were additionally supplemented with 100 units/ml penicillin, 100 μg/ml streptomycin (Corning), and 25 ng/ml amphotericin B (Corning). All cells were maintained at 37°C with 5% CO_2_. Baby hamster kidney cells expressing T7 RNA polymerase under control of a cytomegalovirus promoter (BHK-T7) (56) were maintained in Dulbecco’s minimum essential medium (DMEM; Corning) supplemented to contain 5% FBS (Gibco), with 1 mg/ml Geneticin (Gibco) added during alternate passages. MA104 cells were grown in minimum essential medium with Earle’s salts (EMEM; Corning) supplemented to contain 5% FBS.

### Viruses

Laboratory stocks of reovirus strains rsT1L and rsT3D^I^ were engineered using plasmid-based reverse genetics (19, 56). rsT3D^I^ differs from rsT3D in that it contains an engineered T249I mutation that prevents proteolytic cleavage of attachment protein σ1 (19). The T249I mutation was introduced in the pBacT7-S1T3D plasmid by ‘round the horn PCR (57) with mutagenic primers. Monolayers of BHK-T7 cells at ~50% confluency in 6-well plates were co-transfected using TransIT-LT1 transfection reagent (Mirus Bio LLC) with 0.8 μg of each of ten plasmid constructs representing the T1L or T3D^I^ reovirus genome. After five days of incubation, cells were lysed by two rounds of freezing and thawing, and rsT1L or rsT3D^I^ in lysates was amplified once in L cells prior to use in serial passage experiments. Viral titers were determined by plaque assay using L cells, as described (58).

A laboratory stock of rsSA11 was engineered using reverse genetics (59). Confluent monolayers of BHK-T7 cells in 12-well plates were co-transfected using TransIT-LT1 transfection reagent (Mirus Bio LLC) with 0.27 μg of each of eleven plasmid constructs representing the SA11 rotavirus genome and of each of two plasmids encoding vaccinia virus mRNA capping enzymes, and 0.005 μg of a plasmid encoding the Nelson Bay virus fusion-associated small transmembrane protein. After one day of incubation, supernatants were removed and replaced with serum-free DMEM. After one additional day of incubation, supernatants were removed, and 3.3 × 10^4^ MA104 cells in serum-free DMEM containing 0.5 μg/ml of trypsin were added to each well, followed by incubation at 37°C for three more days. Cells were then lysed by three rounds of freezing and thawing, and rsSA11 in lysates was amplified once in MA104 cells prior to use in serial passage experiments. Virus titer was determined by plaque assay using MA104 cells, as described (60).

### Reovirus serial passage in L cells

For P1, 5 × 10^7^ L cells, which support vigorous reovirus replication, were pelleted and adsorbed with rsT1L or rsT3D^I^ in 10 ml JMEM at an MOI of 1 PFU/cell for 1 h at 37°C. Cell-virus mixtures were transferred to 1 L bottles containing a magnetic stirring rod, and JMEM was added to a total volume of 100 ml. Spinner bottles were incubated at 37°C with stirring for 48 h prior to two rounds of freezing and thawing and removal of large cellular debris by centrifugation at 1000 × g for 5 min. For subsequent passages (P2-P10), an identical protocol was followed, except that 5 × 10^7^ L cells were pelleted and resuspended in an adsorption inoculum of 10 ml of cleared lysate from the previous passage. For each virus, passages were conducted in triplicate lineages, which were maintained throughout the series. Virus titer in each passage for each lineage was determined by plaque assay (58).

### Rotavirus serial passage in MA104 cells

For P1, rsSA11 was activated by incubation with 10 μg/ml of trypsin for 1 h at 37°C then diluted to a final concentration of < 2 μg/ml of trypsin. A confluent monolayer of MA104 cells (~1.5 × 10^7^), which support vigorous rotavirus replication, in a T150 flask was washed with serum-free EMEM adsorbed with rsSA11 in 3 ml EMEM at an MOI of 0.25 PFU/cell for 1 h at 37°C. The inoculum was removed, and the monolayer was washed prior to the addition of 20 ml serum-free EMEM containing 0.5 μg/ml of trypsin. Cells were incubated at 37°C for 48 h prior to two rounds of freezing and thawing and removal of large cellular debris by centrifugation at 1000 × g for 5 min. For subsequent passages (P2-P10), an identical protocol was followed, except that 3 ml of cleared rsSA11 lysate from the previous serial passage was activated by incubation with 1 μg/ml of trypsin for 1 h at 37°C, then used as the adsorption inoculum for the subsequent passage. Passages were conducted in triplicate lineages, which were maintained throughout the series. Virus titer in each passage for each lineage was determined by fluorescent focus assay (FFA).

### Rotavirus fluorescent focus assay

MA104 cells were seeded into 96-well, black-walled plates to achieve a density of ~ 2.25 × 10^4^. Rotavirus stocks were activated by incubation with 10 μg/ml trypsin for 1 h at 37°C then diluted serially in serum-free EMEM. Cells were washed twice with serum-free EMEM then adsorbed with serial dilutions of rotavirus at 37°C for 1 h. Inocula were removed, and cells were washed and incubated in fresh medium at 37°C for 12-18 h. Cells were then fixed with cold methanol, and rotavirus proteins were detected by incubation with polyclonal rotavirus antiserum (Invitrogen) at a 1:1,000 dilution in PBS containing 0.5% Triton X-100 at 37°C, followed by incubation with Alexa Fluor 488-labeled secondary IgG (Invitrogen) and DAPI. Images were captured for four fields of view per well using an ImageXpress Micro XL automated microscope imager (Molecular Devices). Total and infected cells were quantified using MetaXpress high-content image acquisition and analysis software (Molecular Devices). Fluorescent focus units per milliliter of virus stock were calculated based on the surface area of the well quantified and virus inoculum volume and dilution.

### RT-PCR and electrophoretic analysis of DVG profiles

RNA was isolated from cleared serial passage lysates using TRIzol LS Reagent (Invitrogen), according to the manufacturer protocol. Gene segments and DVGs were amplified using a OneStep RT-PCR kit (Qiagen), using the isolated serial passage RNA as a template and gene-specific primers that bind the 5′ and 3′ RNA termini (Table 1). Reaction and thermal cycler conditions were as described by the manufacturer. An extension time of 1 min was used for gene segments < 2 kb, and 3 min was used for segments > 2 kb. RT-PCR products were resolved by electrophoresis in 1.2% agarose in the presence of ethidium bromide and imaged with a VWR photo imager and variable intensity UV transilluminator. RT-PCR product sizes were analyzed in comparison to migration of markers in 100 bp and 1 kb ladders (New England Biolabs).

### DVG sequencing

Selected DNA fragments were excised from agarose gels, purified using a QIAquick Gel Extraction Kit (Qiagen), and sequenced using the Sanger method (Genewiz) and the same primers used for RT-PCR amplification. Sequences were aligned to the reference viral genome using CLC Sequence Viewer 8 (Qiagen).

### rsT1L purification, library preparation, and next-generation sequencing

In two independent preparations, L cells (2 × 10^8^) were pelleted and adsorbed with laboratory stocks of rsT1L reovirus in 20 ml JMEM at an MOI of 10 PFU/cell for 1 h at 37°C. Cell-virus mixtures were transferred to 1 L bottles containing a magnetic stirring rod, and JMEM was added to a total volume of 400 ml. Spinner bottles were incubated at 37°C with stirring for 48 h, then cells were pelleted by centrifugation at 3,000 rpm for 10 min at 4°C. Cell pellets were resuspended in homogenization buffer (25 mM NaCl, 10 mM Tris pH 7.4, 10 mM β-mercaptoethanol) and frozen at −80°C. Cell pellets were thawed and incubated with 0.14% (v/v) deoxycholate for 30 min on ice prior to addition of Vertrel XF and sonication. Virus particles were concentrated by centrifugation at 25,000 × *g* in a 1.2-1.4 g/cm^3^ cesium chloride density gradient. The reovirus-containing fraction was collected and dialyzed at 4°C in virion storage buffer (150 mM NaCl, 15 mM MgCl, 10 mM Tris-HCl, pH 7.4). Purified rsT1L particles were treated with a final concentration of 1 U/μL benzonase (Millipore) for 1h at 37°C to degrade extra-particle nucleic acids. RNA was isolated from benzonase-treated particles using TRIzol LS Reagent (Invitrogen), according to manufacturer’s protocol.

RNA libraries were prepared from RNA isolated from cesium chloride gradient-purified, benzonase-treated rsT1L virions. Contaminating DNA was degraded by treatment with RNase-free DNase I (New England Biolabs) for 10 min at 37°C. RNA was isolated by using TRIzol LS Reagent (Invitrogen), according to the manufacturer’s protocol, and the concentration and quality of RNA was quantified using a Bioanalyzer (Agilent). Library preparation for Illumina sequencing was performed using the NEBNext Ultra II RNA Library Prep Kit for Illumina (New England Biolabs) and a minimum of 5 ng of RNA, according to manufacturer’s instructions. Briefly, RNA was fragmented prior to first-strand and second-strand synthesis and RNAClean XP (Beckman Coulter) purification. PCR enrichment of adaptor-ligated DNA was performed using NEBNext Multiplex Oligos for Illumina (New England Biolabs) to produce Illumina-ready libraries. Libraries were sequenced by 150 base pair paired-end sequencing on a NovaSeq 6000 Sequencing System (Illumina). Assistance with quality control and next-generation sequencing was provided by the Vanderbilt Technologies for Advanced Genomics (VANTAGE) research core.

### Illumina RNA-seq processing and alignment

Raw reads were processed by first removing the Illumina TruSeq adapter using Trimmomatic (61) default settings (command line parameters java -jar trimmomatic.jar PE sample_R1.fastq.gz sample_R2.fastq.gz output_paired_R1.fastq output_unpaired_R1.fastq output_paired_R2.fastq output_unpaired_R2_unpaired.fastq ILLUMINACLIP:TruSeq3-PE.fa:2:30:10 LEADING:3 TRAILING:3 SLIDINGIWINDOW:4:15 MINLEN:36). Reads shorter than 36 bp were removed and low-quality bases (Q score < 30) were trimmed from read ends. The raw FASTQ files were aligned to the reovirus gene segments (NCBI accession numbers M24734.1, AF378003.1, AF129820.1, AF461682.1, AF490617.1, AF174382.1, M14779.1, L19774.1, M14325.1, M13139.1) using the Python2 script *ViReMa* (*Vi*ral Recombination *Ma*pper, version 0.15) (42) using the command line parameters python2 ViReMa.py reference_index input.fastq output.sam --OuputDir sample_virema/ --OutputTag sample_virema -BED --MicroIndelLength 5. The sequence alignment map (SAM) file was processed using the samtools (62) suite to calculate nucleotide depth at each position in a sorted binary alignment map (BAM) file (using command line parameters samtools depth -a -m 0 sample_virema.sorted.bam > sample_virema.coverage).

### Recombination junction analysis

Recombination junctions were first filtered in the forward (5’ → 3’) direction using the *dpylr* package (RStudio). The frequency of each junction was calculated by comparing the depth of the unique junction to the total number of nucleotides in all detected junctions in a library. Junctions were plotted according to the genomic position and colored according to log10 of the frequency using *ggplot2* in RStudio. Recombination frequency was calculated at each genomic position by dividing the number of nucleotides in any junction mapping to the position divided by the total number of nucleotides sequenced at the position.

### Nucleotide composition analysis and sequence microhomology quantification

Nucleotide composition at each position surrounding DVG junctions was determined. To avoid bias of highly replicated DVGs and to more closely reflect the stochastic nature of RNA recombination, each unique detected junction was counted equally rather than weighting by read count (52). Sequences were extracted from a sorted BED file listing the junctions using Rec_Site_Extraction.py with a 30-base pair window. Start site and stop site sequences were separated in Microsoft Excel and the nucleotide frequency at each position was calculated using the *Biostrings* (63) package in RStudio (63). Length of microhomology at junction sites were extracted from *ViReMa* SAM file using the Compiler_Module.py of *ViReMa* and -FuzzEntry --Defuzz 0 flags. The frequency of overlaps ranging from 0 – 10 bp was calculated and compared to an expected probability distribution using uHomology.py.

For junction analyses shown in Tables 2 and 4, regions of sequence similarity were defined as stretches of at least four sequential, identical nucleotides located in the same position or removed by a single nucleotide, relative to the junction site. A single mismatch, shown in black, was tolerated in identical sequences longer than four nucleotides, if at least two additional identical nucleotides were adjacent to it.

### Data Availability

Data generated from Illumina RNA-seq can be accessed at NCBI Sequence Read Archive (SRA) under the BioProject accession PRJNA669717. Code utilized in this report can be accessed at https://github.com/DenisonLabVU/rna-seq-pipeline.

## ACKNOWLEDGEMENTS

We thank the staff at VANTAGE for assistance with next-generation sequencing and Dr. James Chappell for critical reading of the manuscript.

This research was supported in part by the National Center for Research Resources, Grant UL1 RR024975-01, and is now at the National Center for Advancing Translational Sciences, Grant 2 UL1 TR000445-06. The content is solely the responsibility of the authors and does not necessarily represent the official views of the National Institutes of Health.

## Notes

### Competing Interest Statement

The authors have declared no competing interest.

### Summary of Updates

Author name corrected

## REFERENCES

1. Alnaji FG, Brooke CB. 2020. Influenza virus DI particles: Defective interfering or delightfully interesting? PLoS Pathog 16:e1008436.

2. Vignuzzi M, Lopez CB. 2019. Defective viral genomes are key drivers of the virus-host interaction. Nat Microbiol 4:1075–1087.

3. Poirier EZ, Vignuzzi M. 2017. Virus population dynamics during infection. Curr Opin Virol 23:82–87.

4. Rezelj VV, Levi LI, Vignuzzi M. 2018. The defective component of viral populations. Curr Opin Virol 33:74–80.

5. Collaborators GBDCM. 2016. Global, regional, national, and selected subnational levels of stillbirths, neonatal, infant, and under-5 mortality, 1980-2015: a systematic analysis for the Global Burden of Disease Study 2015. Lancet 388:1725–1774.

6. Bouziat R, Hinterleitner R, Brown JJ, Stencel-Baerenwald JE, Ikizler M, Mayassi T, Meisel M, Kim SM, Discepolo V, Pruijssers AJ, Ernest JD, Iskarpatyoti JA, Costes LM, Lawrence I, Palanski BA, Varma M, Zurenski MA, Khomandiak S, McAllister N, Aravamudhan P, Boehme KW, Hu F, Samsom JN, Reinecker HC, Kupfer SS, Guandalini S, Semrad CE, Abadie V, Khosla C, Barreiro LB, Xavier RJ, Ng A, Dermody TS, Jabri B. 2017. Reovirus infection triggers inflammatory responses to dietary antigens and development of celiac disease. Science 356:44–50.

7. Attoui H, Becnel J, Belaganahalli S, Bergoin M, Brussaard CP, Chappell JD, Ciarlet M, del Vas M, Dermody TS, Dormitzer PR, Duncan R, Fang Q, Graham R, Guglielmi KM, Harding RM, Hillman B, Makkay A, Marzachì C, Matthijnssens J, Mertens PPC, Milne RG, Mohd Jaafar F, Mori H, Noordeloos AA, Omura T, Patton JT, Rao S, Maan M, Stoltz D, Suzuki N, Upadhyaya NM, Wei C, Zhou H. 2012. Part II: The Viruses – The Double Stranded RNA Viruses - Family Reoviridae p541 - 637. *In* King AMQ, Adams MJ, Carstens EB, Lefkowitz EJ (ed), Virus taxonomy: classification and nomenclature: Ninth Report of the International Committee on Taxonomy of Viruses. Elsevier Academic Press, San Diego.

8. Guglielmi KM, McDonald SM, Patton JT. 2010. Mechanism of intraparticle synthesis of the rotavirus double-stranded RNA genome. J Biol Chem 285:18123–18128.

9. McDonald SM, Patton JT. 2011. Assortment and packaging of the segmented rotavirus genome. Trends Microbiol 19:136–144.

10. Roy P. 2017. Bluetongue virus structure and assembly. Curr Opin Virol 24:115–123.

11. Trask SD, McDonald SM, Patton JT. 2012. Structural insights into the coupling of virion assembly and rotavirus replication. Nat Rev Microbiol 10:165–177.

12. Dermody TS, Parker JS, Sherry B. 2013. Orthoreoviruses, p 1304–1346. *In* Knipe DM, Howley PM (ed), Fields Virology, Sixth ed, vol 2. Lippincott Williams & Wilkins, Philadelphia.

13. Estes MK, Greenberg HB. 2013. Rotaviruses, p 1347–1401. *In* Knipe DM, Howley PM (ed), Fields Virology, Sixth ed, vol 2. Lippincott Williams & Wilkins, Philadelphia.

14. Biswas S, Li W, Manktelow E, Lever J, Easton LE, Lukavsky PJ, Desselberger U, Lever AM. 2014. Physicochemical analysis of rotavirus segment 11 supports a ‘modified panhandle’ structure and not the predicted alternative tRNA-like structure (TRLS). Arch Virol 159:235–248.

15. Chapell JD, Goral MI, Rodgers SE, dePamphilis CW, Dermody TS. 1994. Sequence diversity within the reovirus S2 gene: reovirus genes reassort in nature, and their termini are predicted to form a panhandle motif. J Virol 68:750–756.

16. Li W, Manktelow E, von Kirchbach JC, Gog JR, Desselberger U, Lever AM. 2010. Genomic analysis of codon, sequence and structural conservation with selective biochemical-structure mapping reveals highly conserved and dynamic structures in rotavirus RNAs with potential cis-acting functions. Nucleic Acids Res 38:7718–7735.

17. Matsuo E, Roy P. 2009. Bluetongue virus VP6 acts early in the replication cycle and can form the basis of chimeric virus formation. J Virol 83:8842–8848.

18. Demidenko AA, Blattman JN, Blattman NN, Greenberg PD, Nibert ML. 2013. Engineering recombinant reoviruses with tandem repeats and a tetravirus 2A-like element for exogenous polypeptide expression. Proc Natl Acad Sci U S A 110:E1867–1876.

19. Kobayashi T, Antar AA, Boehme KW, Danthi P, Eby EA, Guglielmi KM, Holm GH, Johnson EM, Maginnis MS, Naik S, Skelton WB, Wetzel JD, Wilson GJ, Chappell JD, Dermody TS. 2007. A plasmid-based reverse genetics system for animal double-stranded RNA viruses. Cell Host Microbe 1:147–157.

20. Roner MR, Joklik WK. 2001. Reovirus reverse genetics: Incorporation of the CAT gene into the reovirus genome. Proc Natl Acad Sci U S A 98:8036–8041.

21. Zou S, Brown EG. 1992. Identification of sequence elements containing signals for replication and encapsidation of the reovirus M1 genome segment. Virology 186:377–388.

22. AlShaikhahmed K, Leonov G, Sung PY, Bingham RJ, Twarock R, Roy P. 2018. Dynamic network approach for the modelling of genomic sub-complexes in multi-segmented viruses. Nucleic Acids Res 46:12087–12098.

23. Fajardo T, Jr., AlShaikhahmed K, Roy P. 2016. Generation of infectious RNA complexes in Orbiviruses: RNA-RNA interactions of genomic segments. Oncotarget 7:72559–72570.

24. Fajardo T, Jr., Sung PY, Roy P. 2015. Disruption of Specific RNA-RNA Interactions in a Double-Stranded RNA Virus Inhibits Genome Packaging and Virus Infectivity. PLoS Pathog 11:e1005321.

25. Boyce M, McCrae MA, Boyce P, Kim JT. 2016. Inter-segment complementarity in orbiviruses: a driver for co-ordinated genome packaging in the Reoviridae? J Gen Virol 97:1145–1157.

26. Sung PY, Roy P. 2014. Sequential packaging of RNA genomic segments during the assembly of Bluetongue virus. Nucleic Acids Res 42:13824–13838.

27. Ahmed R, Graham AF. 1977. Persistent infections in L cells with temperature-sensitive mutants of reovirus. J Virol 23:250–262.

28. Alam MM, Kobayashi N, Ishino M, Nagashima S, Paul SK, Chawla-Sarkar M, Krishnan T, Naik TN. 2008. Identical rearrangement of NSP3 genes found in three independently isolated virus clones derived from mixed infection and multiple passages of Rotaviruses. Arch Virol 153:555–559.

29. Eaton BT, Gould AR. 1987. Isolation and characterization of orbivirus genotypic variants. Virus Res 6:363–382.

30. Gault E, Schnepf N, Poncet D, Servant A, Teran S, Garbarg-Chenon A. 2001. A human rotavirus with rearranged genes 7 and 11 encodes a modified NSP3 protein and suggests an additional mechanism for gene rearrangement. J Virol 75:7305–7314.

31. Kojima K, Taniguchi K, Kawagishi-Kobayashi M, Matsuno S, Urasawa S. 2000. Rearrangement generated in double genes, NSP1 and NSP3, of viable progenies from a human rotavirus strain. Virus Res 67:163–171.

32. Ramig RF, Samal SK, McConnell S. 1985. Genome RNAS of virulent and attenuated strains of bluetongue virus serotypes 10, 11, 13 and 17. Prog Clin Biol Res 178:389–396.

33. Schnepf N, Deback C, Dehee A, Gault E, Parez N, Garbarg-Chenon A. 2008. Rearrangements of rotavirus genomic segment 11 are generated during acute infection of immunocompetent children and do not occur at random. J Virol 82:3689–3696.

34. Spandidos DA, Graham AF. 1976. Generation of defective virus after infection of newborn rats with reovirus. J Virol 20:234–247.

35. Desselberger U. 1996. Genome rearrangements of rotaviruses. Adv Virus Res 46:69–95.

36. Ni Y, Kemp MC. 1994. Subgenomic S1 segments are packaged by avian reovirus defective interfering particles having an S1 segment deletion. Virus Res 32:329–342.

37. Nonoyama M, Watanabe Y, Graham AF. 1970. Defective virions of reovirus. J Virol 6:226–236.

38. Berger AK, Danthi P. 2013. Reovirus activates a caspase-independent cell death pathway. MBio 4:e00178–00113.

39. Tyler KL, Squier MK, Rodgers SE, Schneider BE, Oberhaus SM, Grdina TA, Cohen JJ, Dermody TS. 1995. Differences in the capacity of reovirus strains to induce apoptosis are determined by the viral attachment protein sigma 1. J Virol 69:6972–6979.

40. Castilho JG, Botelho MV, Lauretti F, Taniwaki N, Linhares RE, Nozawa C. 2004. The in vitro cytopathology of a porcine and the simian (SA-11) strains of rotavirus. Mem Inst Oswaldo Cruz 99:313–317.

41. Bangham CR, Kirkwood TB. 1993. Defective interfering particles and virus evolution. Trends Microbiol 1:260–264.

42. Routh A, Johnson JE. 2014. Discovery of functional genomic motifs in viruses with ViReMa-a Virus Recombination Mapper-for analysis of next-generation sequencing data. Nucleic Acids Res 42:e11.

43. Gribble J, Pruijssers A, Agostini M, Anderson-Daniels J, Chappell J, Lu X, Stevens L, Routh A, Denison M. 2020. The coronavirus proofreading exoribonuclease mediates extensive viral recombination. bioRxiv doi:https://doi.org/10.1101/2020.04.23.057786.

44. Peccoud J, Lequime S, Moltini-Conclois I, Giraud I, Lambrechts L, Gilbert C. 2018. A Survey of Virus Recombination Uncovers Canonical Features of Artificial Chimeras Generated During Deep Sequencing Library Preparation. G3 (Bethesda) 8:1129–1138.

45. Hundley F, Biryahwaho B, Gow M, Desselberger U. 1985. Genome rearrangements of bovine rotavirus after serial passage at high multiplicity of infection. Virology 143:88–103.

46. Tapia F, Laske T, Wasik MA, Rammhold M, Genzel Y, Reichl U. 2019. Production of Defective Interfering Particles of Influenza A Virus in Parallel Continuous Cultures at Two Residence Times-Insights From qPCR Measurements and Viral Dynamics Modeling. Front Bioeng Biotechnol 7:275.

47. Portner A, Kingsbury DW. 1971. Homologous interference by incomplete Sendai virus particles: changes in virus-specific ribonucleic acid synthesis. J Virol 8:388–394.

48. Fodor E, Mingay LJ, Crow M, Deng T, Brownlee GG. 2003. A single amino acid mutation in the PA subunit of the influenza virus RNA polymerase promotes the generation of defective interfering RNAs. J Virol 77:5017–5020.

49. Vasilijevic J, Zamarreno N, Oliveros JC, Rodriguez-Frandsen A, Gomez G, Rodriguez G, Perez-Ruiz M, Rey S, Barba I, Pozo F, Casas I, Nieto A, Falcon A. 2017. Reduced accumulation of defective viral genomes contributes to severe outcome in influenza virus infected patients. PLoS Pathog 13:e1006650.

50. Ooms LS, Kobayashi T, Dermody TS, Chappell JD. 2010. A post-entry step in the mammalian orthoreovirus replication cycle is a determinant of cell tropism. J Biol Chem 285:41604–41613.

51. Yin P, Cheang M, Coombs KM. 1996. The M1 gene is associated with differences in the temperature optimum of the transcriptase activity in reovirus core particles. J Virol 70:1223–1227.

52. Jaworski E, Routh A. 2017. Parallel ClickSeq and Nanopore sequencing elucidates the rapid evolution of defective-interfering RNAs in Flock House virus. PLoS Pathog 13:e1006365.

53. Lazzarini RA, Keene JD, Schubert M. 1981. The origins of defective interfering particles of the negative-strand RNA viruses. Cell 26:145–154.

54. Lui WY, Yuen CK, Li C, Wong WM, Lui PY, Lin CH, Chan KH, Zhao H, Chen H, To KKW, Zhang AJX, Yuen KY, Kok KH. 2019. SMRT sequencing revealed the diversity and characteristics of defective interfering RNAs in influenza A (H7N9) virus infection. Emerg Microbes Infect 8:662–674.

55. Saira K, Lin X, DePasse JV, Halpin R, Twaddle A, Stockwell T, Angus B, Cozzi-Lepri A, Delfino M, Dugan V, Dwyer DE, Freiberg M, Horban A, Losso M, Lynfield R, Wentworth DN, Holmes EC, Davey R, Wentworth DE, Ghedin E, Group IFS, Group IFS. 2013. Sequence analysis of in vivo defective interfering-like RNA of influenza A H1N1 pandemic virus. J Virol 87:8064–8074.

56. Kobayashi T, Ooms LS, Ikizler M, Chappell JD, Dermody TS. 2010. An improved reverse genetics system for mammalian orthoreoviruses. Virology 398:194–200.

57. Hemsley A, Arnheim N, Toney MD, Cortopassi G, Galas DJ. 1989. A simple method for site-directed mutagenesis using the polymerase chain reaction. Nucleic Acids Res 17:6545–6551.

58. Berard A, Coombs KM. 2009. Mammalian reoviruses: propagation, quantification, and storage. Curr Protoc Microbiol Chapter 15:Unit15C.11.

59. Kanai Y, Komoto S, Kawagishi T, Nouda R, Nagasawa N, Onishi M, Matsuura Y, Taniguchi K, Kobayashi T. 2017. Entirely plasmid-based reverse genetics system for rotaviruses. Proc Natl Acad Sci U S A 114:2349–2354.

60. Arnold M, Patton JT, McDonald SM. 2009. Culturing, storage, and quantification of rotaviruses. Curr Protoc Microbiol Chapter 15:Unit 15C 13.

61. Bolger AM, Lohse M, Usadel B. 2014. Trimmomatic: a flexible trimmer for Illumina sequence data. Bioinformatics 30:2114–2120.

62. Li H, Handsaker B, Wysoker A, Fennell T, Ruan J, Homer N, Marth G, Abecasis G, Durbin R. 2009. The Sequence Alignment/Map format and SAMtools. Bioinformatics 25:2078–2079.

63. Pagès H, Aboyoun P, Gentleman R, DebRoy S. 2020. Biostrings: Efficient manipulation of biological strings. R package version 2560 [Computer Software].

