## Supplemental Figures for "Reovirus RNA recombination is sequence directed and generates internally deleted defective genome segments during passage"

**T1L L1 DVG**
GCTACACGTTCCACGACACAATGTCATCCATGATACTGACTCAGTTTAGACCGTTCATTGAAAGCATTTCAGGTATCACT
GACCAATCGAATGACGTGTTTGAAGATGCAGCAAAAGCATTCTCTATGTTTACTCGCAGCGACGTCTACAAGGCGCTGGA
TGAAATACCTTTCTCTGATGATGCGATGCTTCCCATCCCTCCAACTATATATACTAAACCATCTCACGATTCATATTATT
ACATTGATGCTCTAAACCGTG/GTGCGTCAACTCATC

**T1L M1 DVG**
GCTATTCGCGGTCATGGCTTACATCGCAGTTCCTGCGGTGGTGGATTCACGTTCAAGTGAGGCTATTGGACTGCTAGAATC
GTTT/ACAATGCGAGACTAGCTTTCCGATCTGACTTGGCGTGATCCGTGACATGCGTAGTGTGACACCTGCCCCTAGGTCA
ATGGGGGTAGGGGGCGGGCTAAGACTACGTACGCGCTTCATC

**T1L S4 DVG**
GCTATTTTTGCCTCTTCCCAAACGTTGTCGCAATGGAGGTGTGCTTGCCCAACGGTCATCAGATCGTGGACTTGATTAAC
AACGCTT/AAGGAACGGCTAAGTTAAAGACAGTGCGCAAGCTAGTGGATTCAGTCAATCATGCGTGGGGCGTCGAGAAGAT
CAGATATGCGCTTGGACCAGGTGGCATGACGGGATGGTACAACAGGACTATGCAGCAGGCCCCCATTGTACTAACTCCCG
CTGCTCTCACAATGTTCTCAGACACCACCAAGTTCGGGGATTTGGATTATCCGGTGATGATTGGCGATCCGATGATTCTT
GGCTAAACACCCCCATCTTCACAGCGCCGGGCCTGACCAACCTGGTGTGACGTGGGACAGGCTCCATTCATC

**T3D$^{I}$ L1 DVG**
GCTACACGTTCCACTTCCACGACAATGTCATCCATGATACTGACTCAGTTTGGACCGTTCATTGAGAGCATT/TCAGGTTC
AACAGCCACCTCTACTGAGCATACTGCTAATAATAGTACGATGATGGAAACTTTCCTGACAGTATGGGGACCCGAACATAC
TGACGACCCTGACGTCTTACGTTTAATGAAGTCTTTAACTATTCAAAGAAATTACGTATGTCAAGGTGATGATGGATTAAT
GATTATCGATGGGACTACTGCTGGT/GCCGGAATGGTAATGACTGCGACTGGAGTTGCTGTCGACATCTATCTGGAGGATA
TACATGGCGGTGGTCGGTCACTTGGACAGAGATTCATGACTTGGATGCGACAGGAAGGACGGTCAGCGTGAGTCTACCATG
GGTCGTGGTGCGTCAACTCATC

**T3D$^{I}$ M1 DVG**
GCTATTCGCGGTCATGGC/ACTGGACGTGGAGCTGCATACAGTGCGAGACTAGCTTTCCGATCTGACTTGGCGTGATCCGT
GACATGCGTAGTGTGACACCTGCTCCTAGGTCAATGGGGGTAGGGGGCGGGCTAAGACTACGTACGCGCTTCATC

**T3D$^{I}$ S4 DVG**
GCTATTTTTGCCTCTTCCCAGACGTTGTCGCAATGGAGGTGTGCTTGCCCAACGGTCATCAGGTCGTGGACTTGATTAAC
AACGCTTTTGAAGGTCGTGTATCAATCTACAGCGCGCAAGAGGGA/TGGGACAGGCTTCATTCATC

**Figure S1. Sequences of reovirus DVGs.** RT-PCR products amplified using primers that bind the 5′ and 3′ termini of the L1, M1, and S4 reovirus segments, smaller than the full-length segments, and indicated in Figure 2 were excised from agarose gels and sequenced. 5′ and 3′ UTRs are colored gray, recombination sites are indicated by a black forward slash, and ORF sequences upstream or downstream of recombination sites are colored in shades of purple (L1), blue (M1), or green (S4).

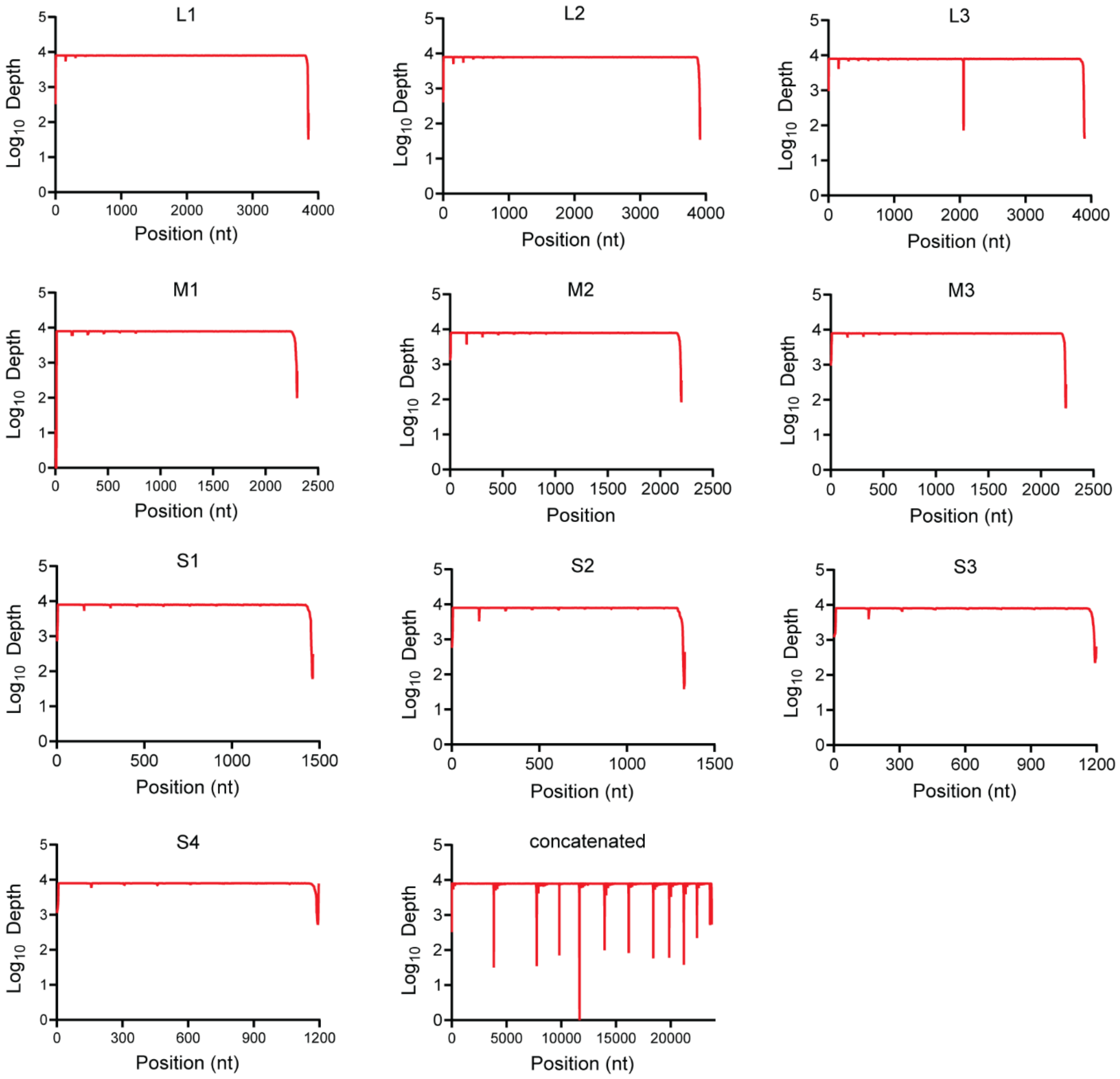


**Figure S2.** rsT1L reovirus sequence depth. Graphs showing sequence depth at each nucleotide position across the indicated rsT1L genome segment or the concatenated reovirus genome (L1, L2, L3, M1, M2, M3, S1, S2, S3, S4). Mean depth for two samples of sequenced virion RNA is shown.

.

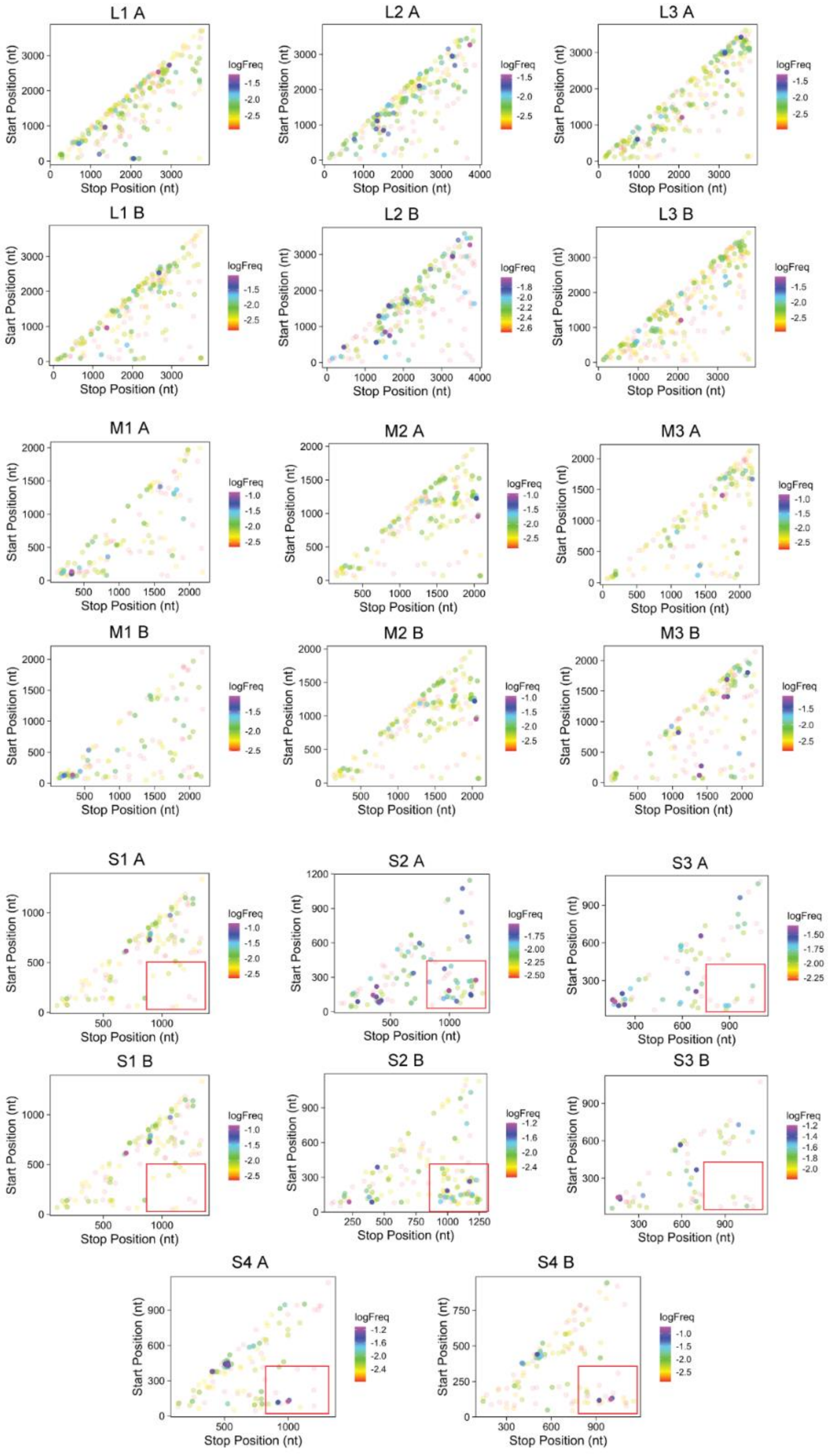
L1 A
L2 A
L3 A
L1 B
L2 B
L3 B
M1 A
M2 A
M3 A
M1 B
M2 B
M3 B
S1 A
S2 A
S3 A
S1 B
S2 B
S3 B
S4 A
S4 B
Start Position (nt)
Stop Position (nt)
logFreq

**Figure S3. rsT1L reovirus recombination junction maps.** Graphs showing recombination junction site location and frequency in sequenced rsT1L virion RNA for each reovirus genome segment. Junction sites are indicated by dots whose position corresponds to upstream and downstream sequences that are merged to form a novel junction. Junction frequency is indicated by dot color, according to the legend to the right of each image. Junction maps for independently purified and sequenced virion RNA preparations are shown in separate images, labeled A or B. Red boxes shown on S4 junction plots outline junctions in which the 5′ and 3′ segment termini are merged.

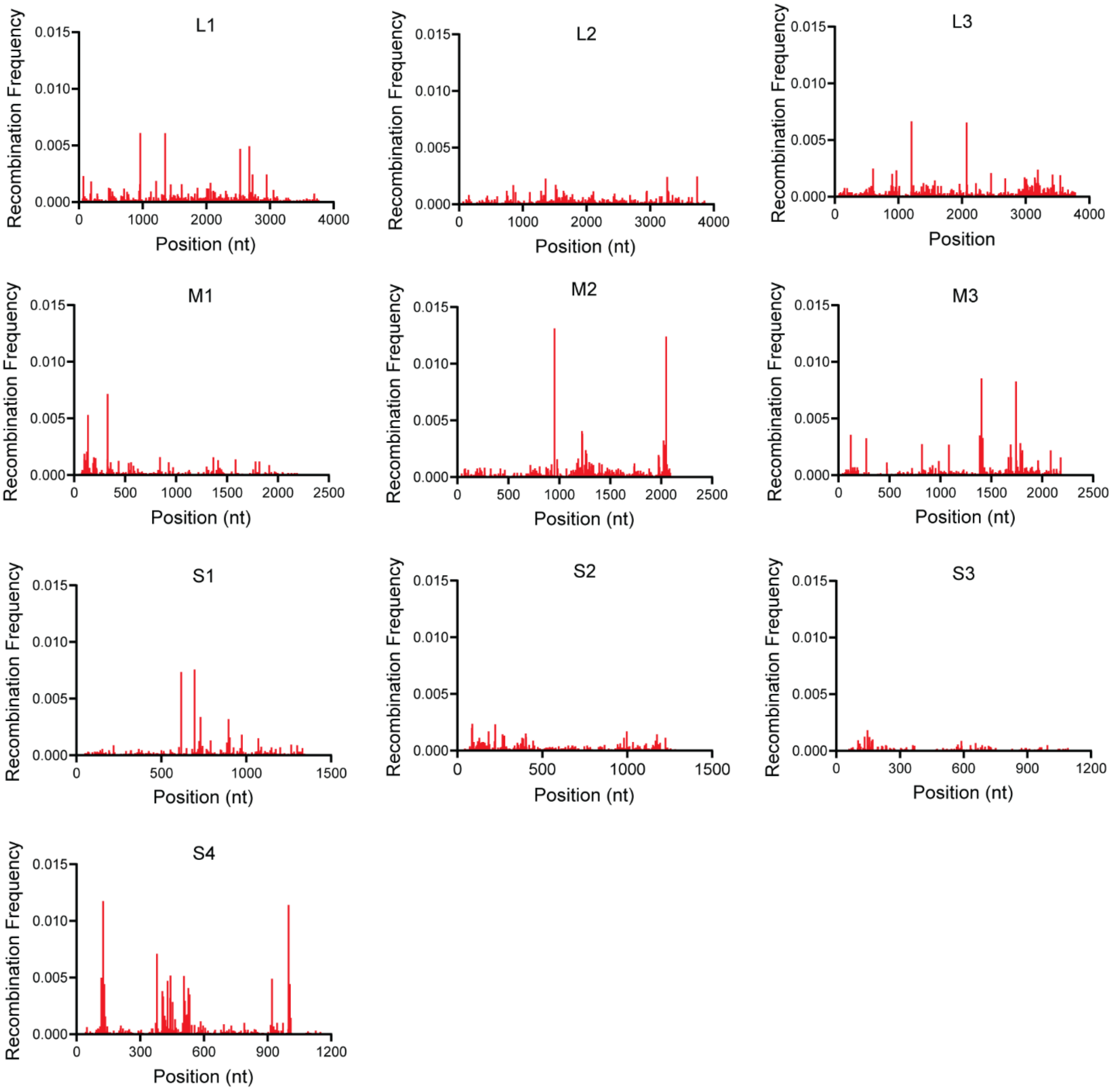


**Figure S4. rsT1L reovirus positional recombination frequency.** Graphs showing recombination frequency at each nucleotide position across the indicated rsT1L genome segment. Mean positional recombination frequency for two samples of sequenced virion RNA is shown.

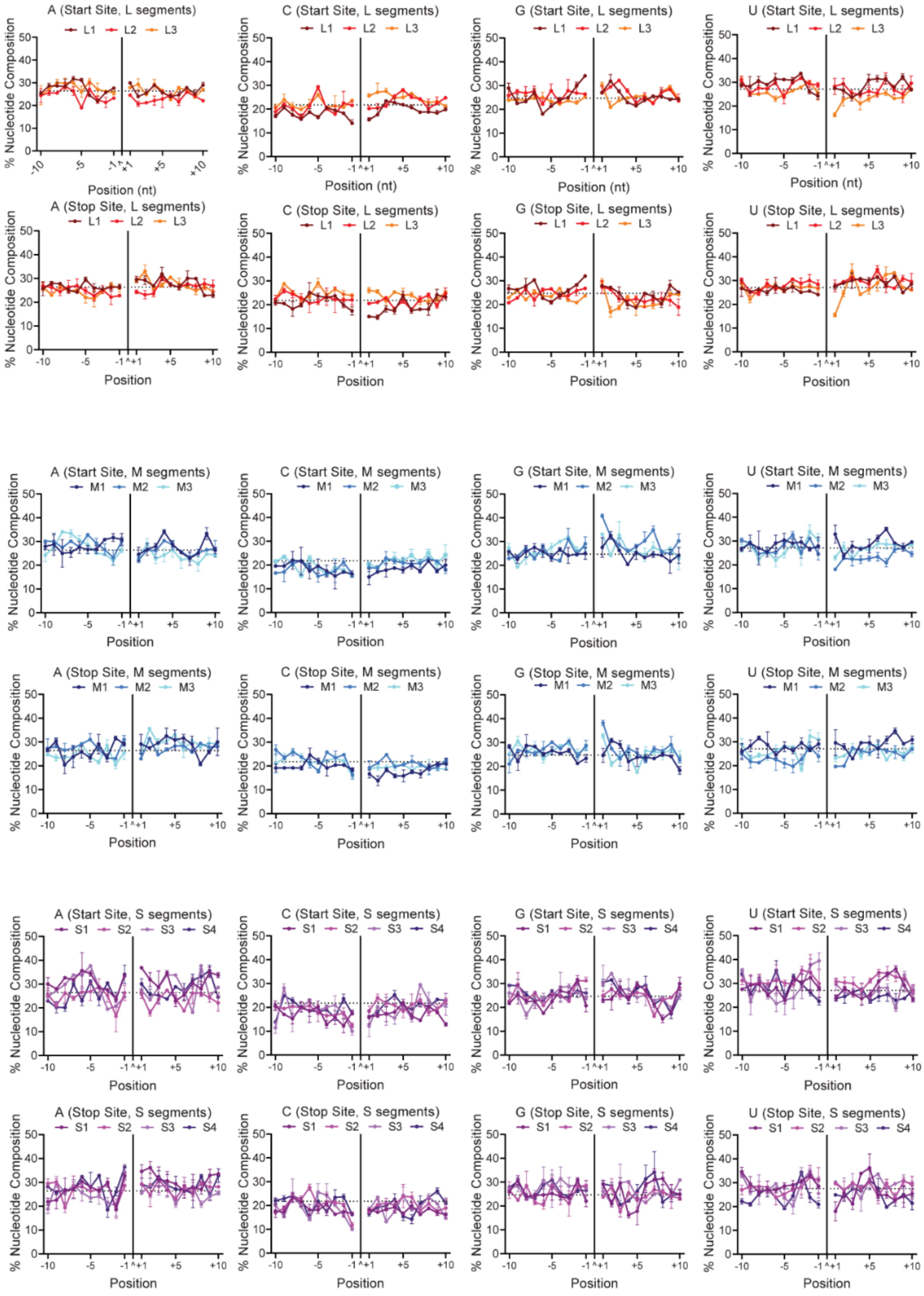
A (Start Site, L segments)
C (Start Site, L segments)
G (Start Site, L segments)
U (Start Site, L segments)
L1 L2 L3
A (Stop Site, L segments)
C (Stop Site, L segments)
G (Stop Site, L segments)
U (Stop Site, L segments)
A (Start Site, M segments)
C (Start Site, M segments)
G (Start Site, M segments)
U (Start Site, M segments)
M1 M2 M3
A (Stop Site, M segments)
C (Stop Site, M segments)
G (Stop Site, M segments)
U (Stop Site, M segments)
A (Start Site, S segments)
C (Start Site, S segments)
G (Start Site, S segments)
U (Start Site, S segments)
S1 S2 S3 S4
A (Stop Site, S segments)
C (Stop Site, S segments)
G (Stop Site, S segments)
U (Stop Site, S segments)
% Nucleotide Composition
Position (nt)
Position
-10 -5 -1 ^+1 +5 +10

**Figure S5. rsT1L reovirus nucleotide frequency at junctions.** Nucleotide composition was calculated as the percent adenosine (A), cytosine (C), guanine (G), and uracil (U) at each position in a 10-base pair region surrounding the DVG start and stop sites for individual L (red/orange), M (blue), and S (purple) segments. The junction is labeled as a caret (^) and denoted with a solid black line. Positions upstream (-10 to -1) and downstream (+1 to +10) of the junction position are indicated. Each point represents a mean (*n* = 2) and error bars represent standard error.
